# Data-driven multiscale model of macaque auditory thalamocortical circuits reproduces in vivo dynamics

**DOI:** 10.1101/2022.02.03.479036

**Authors:** Salvador Dura-Bernal, Erica Y Griffith, Annamaria Barczak, Monica N O’Connell, Tammy McGinnis, Charles E Schroeder, William W Lytton, Peter Lakatos, Samuel A Neymotin

## Abstract

We developed a biophysically-detailed model of the macaque auditory thalamocortical circuits, including primary auditory cortex (A1), medial geniculate body (MGB) and thalamic reticular nuclei (TRN), using the NEURON simulator and NetPyNE multiscale modeling tool. We simulated A1 as a cortical column with a depth of 2000 μm and 200 μm diameter, containing over 12k neurons and 30M synapses. Neuron densities, laminar locations, classes, morphology and biophysics, and connectivity at the long-range, local and dendritic scale were derived from published experimental data. The A1 model included 6 cortical layers and multiple populations of neurons consisting of 4 excitatory and 4 inhibitory types, and was reciprocally connected to the thalamus (MGB and TRN), mimicking anatomical connectivity. MGB included core and matrix thalamocortical neurons with layer-specific projection patterns to A1, and thalamic interneurons projecting locally. Auditory stimulus-related inputs to the MGB were simulated using phenomenological models of the cochlear/auditory nerve and the inferior colliculus. The model generated cell type and layer-specific firing rates consistent with experimentally observed ranges, and accurately simulated the corresponding local field potentials (LFPs), current source density (CSD), and electroencephalogram (EEG) signals. Laminar CSD patterns during spontaneous activity, and in response to speech input, were similar to those recorded experimentally. Physiological oscillations emerged spontaneously across frequency bands without external rhythmic inputs and were comparable to those recorded in vivo. We used the model to unravel the contributions from distinct cell type and layer-specific neuronal populations to oscillation events detected in CSD, and explored how these relate to the population firing patterns. Overall, the computational model provides a quantitative theoretical framework to integrate and interpret a wide range of experimental data in auditory circuits. It also constitutes a powerful tool to evaluate hypotheses and make predictions about the cellular and network mechanisms underlying common experimental measurements, including MUA, LFP and EEG signals.

## 1. Introduction

The auditory system is involved in a number of crucial sensory functions, including speech processing (Hamilton et al. 2021; Matsumoto et al. 2011; Fontolan et al. 2014; Gourévitch et al. 2008), sound localization (Andéol et al. 2011; Carlile, Martin, and McAnally 2005; Ahveninen, Kopčo, and Jääskeläinen 2014), pitch discrimination (Tramo, Shah, and Braida 2002; Tramo et al. 2005; Dykstra et al. 2012; Hyde, Peretz, and Zatorre 2008), and voice recognition (Latinus et al. 2013; Holmes and Johnsrude 2021). Aberrations along this pathway can result in a wide variety of pathologies. Hearing loss, for example, can result from lesions in either the peripheral (Merchant and Rosowski 2008; Raveh et al. 2002) or central (Taniwaki et al. 2000; Brody et al. 2013; Cavinato et al. 2012) parts of this pathway, while other abnormalities can result in increased sensitivity to sound volume (Baguley 2003) or difficulty processing music (Zendel et al. 2015; Albouy et al. 2013).

Achieving a full understanding of this system is complicated by the many interareal pathways, the complexity of the inter- and intralaminar circuitry, the heterogeneity of neuronal cell types and behaviors, and the diversity of network coding mechanisms. A growing body of experimental data, with findings drawn from different methods at different biological scales, begets the need for a framework which can integrate these disparate findings and be used to investigate the system as a whole. The model we present here uses multiscale information with macaque-specific cortical dimensions, a diversity of excitatory and inhibitory cell types with data-driven electrophysiology (Povysheva et al. 2007), data-driven population density and connectivity, detailed thalamic circuits (including a full thalamocortical loop) (Markov et al. 2011), and realistic inputs from upstream structures such as cochlea and inferior colliculus.

Given the multiscale detail of this model, it can also be used to make predictions about the cellular and network-level mechanisms governing oscillatory dynamics in auditory cortex, which is important since cortical oscillations are known to play a prominent role in neural information processing. In auditory cortex, these oscillations may be particularly important for speech processing (Dimitrijevic et al. 2017; Schroeder et al. 2008; Giraud and Poeppel 2012; Ghitza 2011), with oscillations in different frequency bands synchronizing to and tracking the dynamic properties of speech waveforms (Giraud and Poeppel 2012). In some cases, oscillatory behavior in the auditory cortex can even be used to predict speech intelligibility (Ghitza 2011; Dimitrijevic et al. 2017). Abnormalities in auditory cortex oscillations have been observed in pathologies that include auditory processing deficits, such as schizophrenia (Y. Hirano et al. 2020; Spencer et al. 2009; S. Hirano et al. 2018) and autism spectrum disorder (Gandal et al. 2010; De Stefano et al. 2019). Increased oscillatory activity at rest (Y. Hirano et al. 2020), strong cross-frequency synchronization (Spencer et al. 2009; S. Hirano et al. 2018), and impaired phase-locking between auditory cortex oscillations and incoming speech stimuli (Gandal et al. 2010; De Stefano et al. 2019; Jochaut et al. 2015) have been observed in these disorders, and may help explain some of the auditory processing related deficits seen in these disease states (Spencer et al. 2009; Paciello et al. 2021; Lakatos et al. 2019; Edgar et al. 2015; Gandal et al. 2010).

Our model can be used to investigate the cellular-and network-level mechanisms underlying thalamocortical oscillations, by first reproducing the oscillations in silico and then examining the activity which occurs at different scales: from subthreshold currents and dendritic effects, to activity in different thalamic pathways (e.g. core vs. matrix). Bridging these hierarchical levels also allows us to gain insights into the biophysical mechanisms underlying activity observed during experimental recordings that occur at different scales (e.g. single cell recordings, multiunit activity, local field potentials, current source density, electro- and magneto-encephalography).

## 2. Results

### 2.1 Development of a data-driven model of macaque auditory thalamocortical circuits

We developed a biophysically-detailed model of macaque auditory thalamocortical circuits, including medial geniculate body (MGB), thalamic reticular nucleus (TRN) and primary auditory cortex (A1). To provide input to the thalamic populations, we connected a phenomenological model of the cochlear nucleus, auditory nerve and inferior colliculus (IC). This resulted in a realistic model capable of processing arbitrary input sounds along the main stages of the macaque auditory pathway (Fig. 1A). While details of each stage can be found in the Methods section, the current section includes an overall description of the main features of the model.

**Figure 1.**
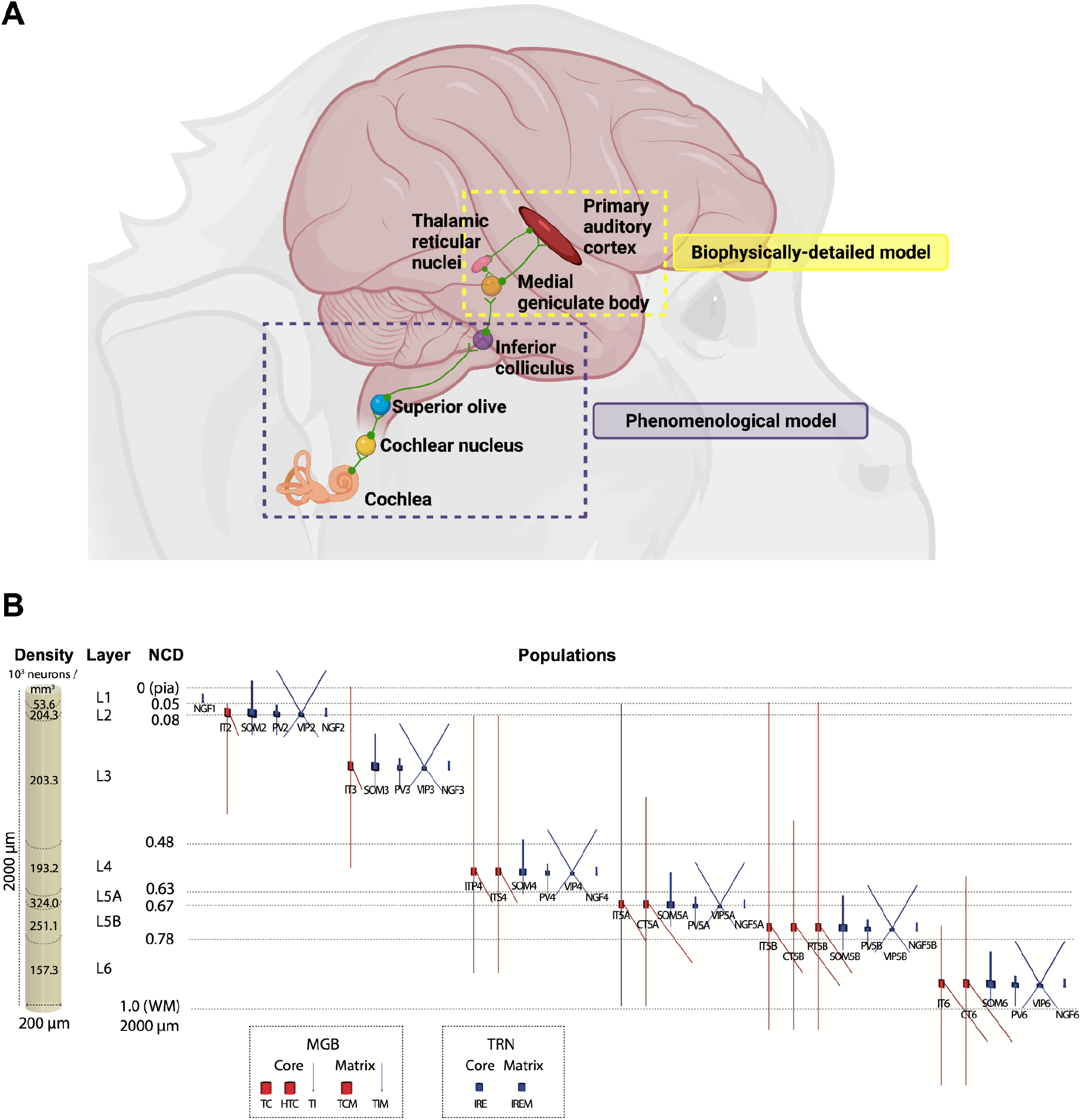
Overview of (A) the macaque auditory system model and (B) the biophysically-detailed auditory thalamocortical circuits model (B). A) A phenomenological model is used to capture the transformation of input sound into electrical impulses in the cochlea, superior olive and inferior colliculus (IC). Output from IC then drives a detailed biophysical model of auditory thalamocortical circuits, including medial geniculate body (MGB), thalamic reticular nuclei (TRN) and primary auditory cortex (A1). Note: many of the connections are bidirectional, but not shown for simplicity. B) Dimensions of simulated A1 column with laminar cell densities, layer boundaries, cell morphologies and distribution of populations. Medial geniculate body (MGB) and thalamic relay nuclei (TRN) populations and simplified morphologies are shown in the bottom, highlighting distinct core- and matrix-projecting populations. All models are conductance-based with multiple ionic channels tuned to reproduce the cell’s electrophysiology. NCD is normalized cortical depth with values ranging from 0 (pia) to 1 (white matter, WM).

We reconstructed a cylindrical volume of 200 um radius and 2000 um depth of A1 tissue (Fig. 1B). The model included 12,187 neurons and over 25 million synapses, corresponding to the full neuronal density of the volume modeled. The model was divided into 7 layers -- L1, L2, L3, L4, L5A, L5B and L6 -- each with boundaries, neuronal densities, and distribution of cell types derived from experimental data (J. A. Winer and Larue 1996; Coen-Cagli, Kanitscheider, and Pouget 2017; Markram et al. 2015; Billeh et al. 2020; Kelly and Hawken 2017; Tremblay, Lee, and Rudy 2016; Lefort et al. 2009; Schuman et al. 2019; Harris and Shepherd 2015; Huang, Larue, and Winer 1999). We included the four main classes of excitatory neurons: intratelencephalic (IT), present in all layers except L1; spiny stellate (ITS) in L4, pyramidal tract (PT) in L5B, and corticothalamic (CT) in L5A, L5B and L6. The dendritic length of cell types in different layers was adapted according to experimental data. While many previous cortical models only include one or two interneuron types, we incorporated a greater diversity of cell type by including four classes of interneurons: somatostatin (SOM), parvalbumin (PV), vasoactive intestinal peptide (VIP), and neurogliaform (NGF). All 4 classes were present in all layers except L1, which only included NGF. The MGB included two types of thalamocortical neurons and thalamic interneurons. The TRN included a population of inhibitory cells. Thalamic populations were in turn divided into core and matrix subpopulations, each with distinct wiring. The total number of thalamic neurons was 721, with cell densities and ratios of the different cell types derived from published studies.

Connectivity in the model was established for each pair of the 42 cortical and thalamic populations resulting in layer- and cell type-specific projections (Fig. 2). Each projection between populations was characterized by a probability of connection and unitary connection strength (in mV), defined as the PSP amplitude in a postsynaptic neuron in response to a single presynaptic spike. The probability of connection from cortical inhibitory populations decayed exponentially with cell-to-cell distance. Synapses were distributed along specific regions of the somatodendritic tree for each projection. Excitatory synapses included colocalized AMPA and NMDA receptors, and inhibitory synapses included different combinations of slow GABA_A_, fast GABA_A_ and GABA_B_ receptors, depending on cell types. The values for all the connectivity parameters were derived from over 30 published experimental studies (see Methods). Afferent projections from other brain regions were modeled by providing background independent Poisson spiking inputs to apical excitatory and basal inhibitory synapses, adjusted for each cell type to result in low spontaneous firing rates (∼1 Hz). Where available, we used data from the NHP auditory system, but otherwise resorted to data from other species, including rodent, cat and human. We employed automated parameter optimization methods to fine tune the connectivity strengths to obtain physiologically constrained firing rates across all populations.

**Figure 2.**
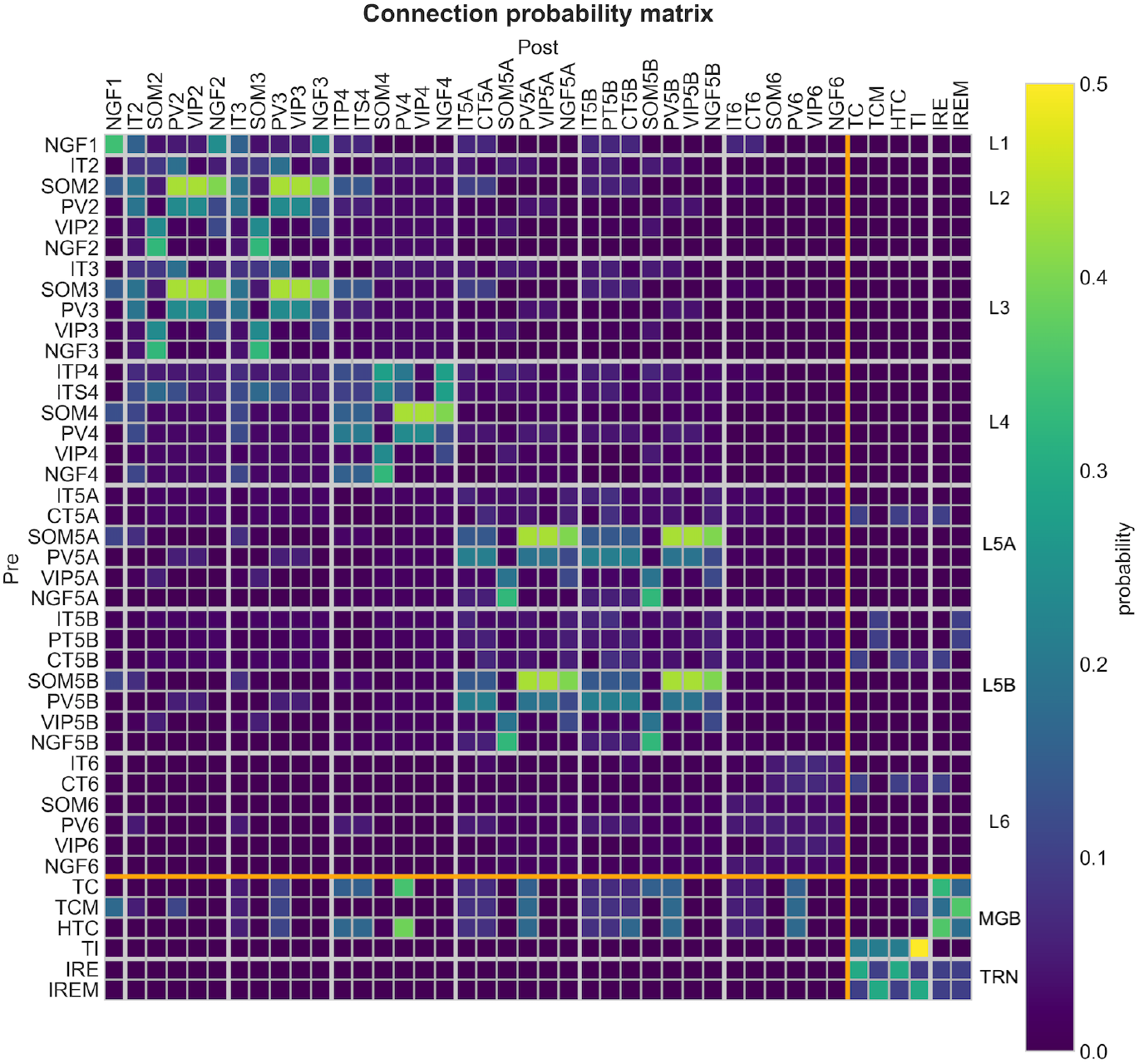
Connectivity matrix of model thalamocortical populations. Probability of connection between all 36 cortical and 6 thalamic populations (thick white lines separate layers and thick orange line separates cortex from thalamus). Note that the gain from IC → MGB is not shown here since there is no feedback connection from MGB → IC in our model, given that we use phenomenological cochlea/IC models (see Methods).

To achieve variability in the baseline model we modified the randomization seeds used to generate the probabilistic connections and spike times of Poisson background inputs. Specifically, we ran 25 simulations with different seeds (5 connectivity x 5 input seeds), each for 11.5-second simulations (first 1.5 seconds required to reach steady state). This resulted in 250 seconds of simulated data, which is comparable to some of the macaque experimental datasets used. Modeling results that include statistics were calculated across all of the 25 × 10 second simulations, which provided a measure of the robustness of the model to variations in connectivity and inputs and is comparable to the variability across different macaques and trials, respectively.

We developed the models using NetPyNE (Dura-Bernal et al. 2019) and parallel NEURON (Lytton et al. 2016). Overall, we ran over 500,000 simulations in order to tune the model parameters and explore model responses to different inputs and conditions. This required over 5 million core hours on several supercomputers, primarily Google Cloud Platform. All model source code, results, and comparisons to experimental data are publicly available on ModelDB and Github.

### 2.2 Cell type and layer-specific activity recorded at multiple scales

The model generated layer and cell type-specific spontaneous activity (Fig. 3). Distinct spiking patterns were recorded across thalamus and cortex (Fig. 3A): TC and TRN showed clear alpha oscillations (∼8 Hz); cortical granular and supragranular layers exhibited a similar oscillatory pattern but more diffuse over time and with higher variability in the peak amplitudes; and infragranular layers showed more tonic firing and a slower delta (∼2 Hz) oscillation. Spiking responses also varied across cell types within a layer, e.g. L5B IT cells fired tonically whereas L5B CT cells showed phasic firing at delta frequency (only 2 peaks of activity). Overall, average spontaneous firing rates were below 5 Hz for excitatory neurons and below 20 Hz for inhibitory neurons, consistent with experimental data (X. Wang et al. 2005; Sakata and Harris 2009; Eggermont 1992; Hromádka, Deweese, and Zador 2008). Spontaneous activity was simulated by driving the thalamic and cortical neurons with non-rhythmic Poisson-distributed low amplitude background inputs. Therefore, the distinct responses of neural populations must be a consequence of their heterogeneous biophysical properties and synaptic connectivity.

**Figure 3.**
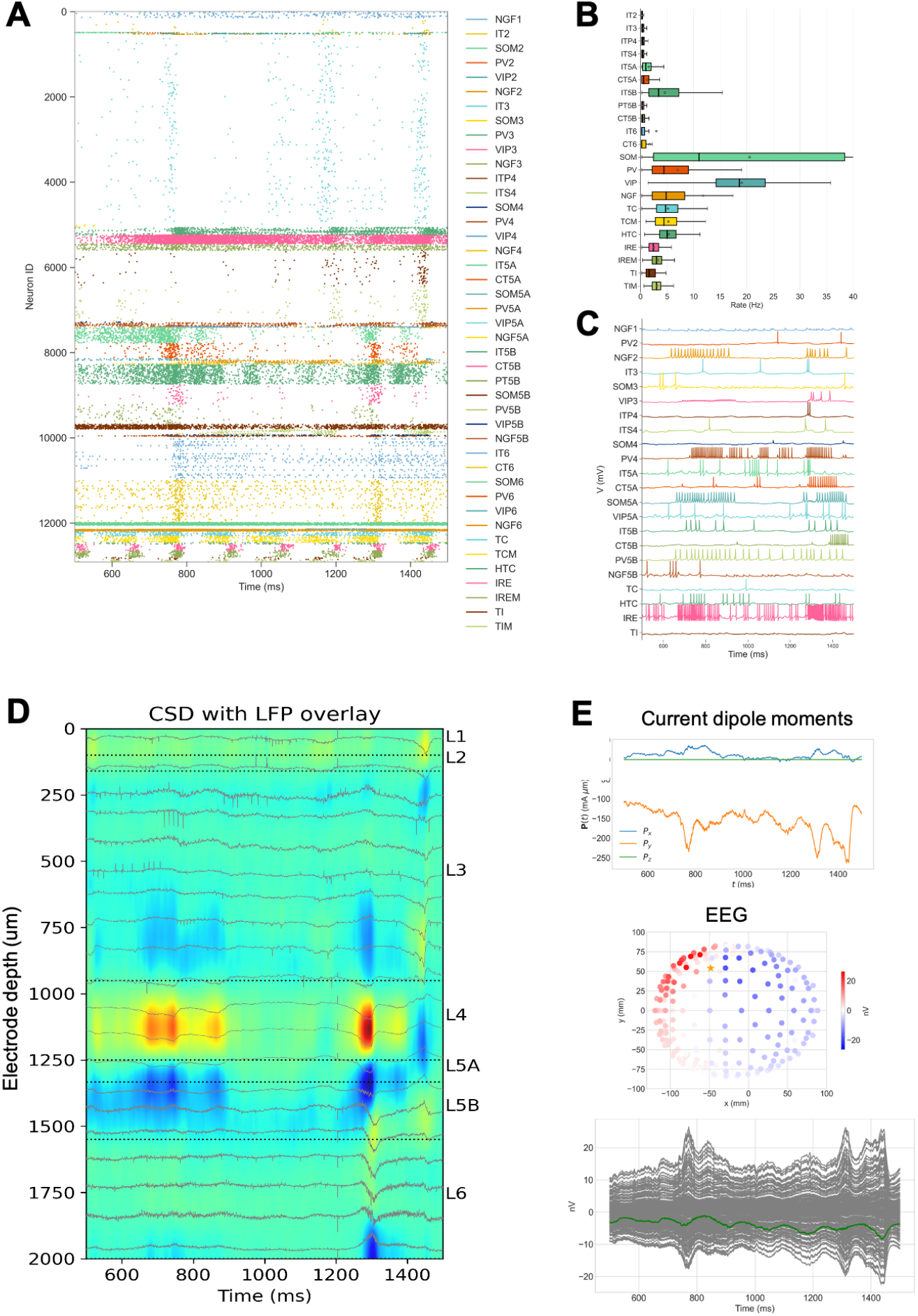
Cell type and layer-specific activity recorded at multiple scales. A) Spiking raster plot. B) Boxplot statistics of the firing rate of all populations (interneurons grouped across all layers). C) Example voltage traces for different cell types and layers. D) Laminar CSD with LFP overlaid. E) Sum of current dipole moments (Px, Py, Pz indicate dipole moment orientations) across all neurons used to calculate EEG signals recorded from scalp electrodes distributed across a volume conduction head model (each electrode in gray; mean in green).

The model responses were recorded and analyzed at multiple scales (Fig. 3): neuronal membrane voltage traces (Fig. 3C), spike times (Fig. 3A), firing rate statistics (Fig. 3B), local field potentials (LFPs) and current source density (CSD) analysis (Fig. 3D), and current dipole moments and electroencephalogram (EEG) signals (Fig. 3E). These measurements represent the same underlying biophysical phenomenon as evidenced by activity features shared across them, e.g. activity peaks around 800 ms and 1300 ms (see Figs. 3A, 3D, 3E). This illustrates how the model can be used to interpret common experimental measurements (MUA, LFP, EEG) and relate them to the underlying biophysical circuit properties. In Section 2.4 we use this approach to disentangle the layer and cell type-specific biophysical sources of an oscillation event.

To examine patterns of LFP/CSD activity in NHP recordings, we first determined the supragranular, granular, and infragranular layer depths for each macaque subject, as done previously (Lakatos et al. 2016). In macaques, the determination of the supragranular, granular, and infragranular layer depths relied on functional demarcation of these regions based on responses to preferred modality stimuli. For each NHP subject, we examined an averaged CSD profile resulting from the presentation of a stimulus which provoked an excitatory response in A1 (e.g. clicks, best frequency tones). An early sink in this CSD profile indicated the presence of the granular layer, while source / sink pairs above and below the granular layer designated the presence of the supragranular and infragranular layers, respectively.

Characteristic laminar LFP/CSD activity patterns recorded experimentally in macaques were qualitatively reproduced in model simulations (Fig. 4). For example, during spontaneous activity we observed examples in both experiment and model showing: 1) ∼50 ms long current sinks (red) in the granular layer together with current sources (blue) immediately above and below, plus current sources (blue) in the most superficial electrodes (Fig. 4A); 2) ∼150 ms long current sinks fluctuating around the border of the granular and infragranular layers with current sources immediately below, and again in the most superficial electrodes (Fig. 4B); and 3) ∼150 ms long current sources in the granular layer with current sinks above and below (Fig. 4C). While reproducing responses to specific speech utterances is outside the scope of this paper, examples of laminar LFP/CSD responses to speech are shown in Fig. 4 and illustrate that the cochlea and IC model can be used to provide auditory stimuli to the biophysical thalamocortical model, which in turn generates activity patterns that resemble those recorded experimentally. For example, ∼150-200ms long current sinks in the granular layer, with alternating current sources and sinks in the infragranular layers (Figs. 4D,E); and short ∼30ms current source in granular layer with similar duration current sources in the supra- and infragranular layers (Fig. 4F). Although there are similarities, these three examples of spontaneous and speech responses also illustrate the variability observed both within and between the experimental and modeling datasets. In the next section, we further quantify this variability in terms of the oscillatory power of spontaneous responses.

**Figure 4.**
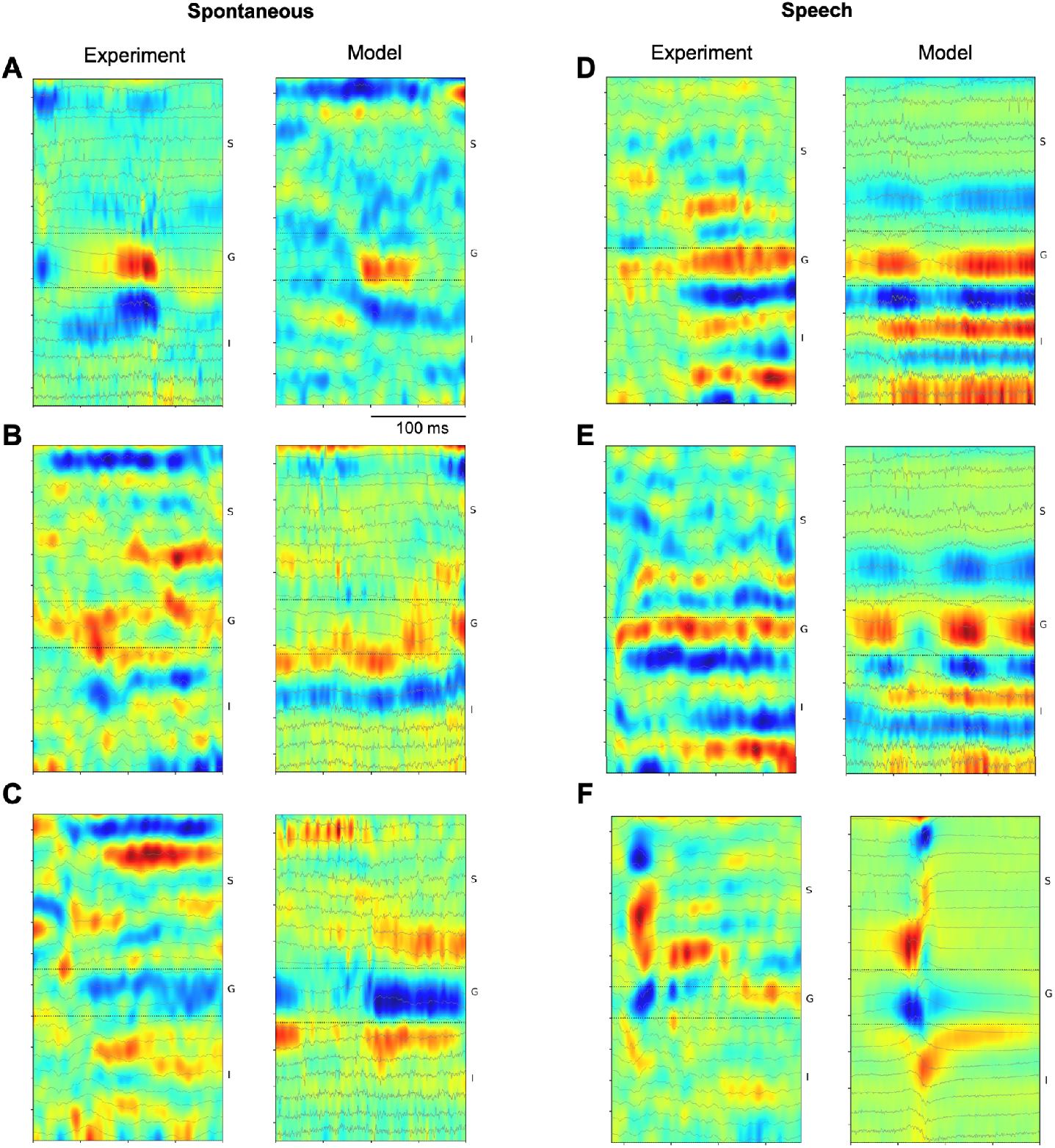
Comparison of example laminar LFP/CSD during spontaneous activity and speech for macaque vs model. These examples illustrate the variability of patterns recorded for each condition, and how the model reproduced some key features of each example pattern (e.g. in top-left example, a 50 ms long current sink in the granular layer and current source infragranular layer). Transmembrane currents (sinks and sources) in CSD color maps are color-coded red and blue, respectively. Y-axis represent LFP and related CSD channels at depths spanning pia to white matter, with supragranular (S), granular (G) and infragranular (I) layers indicated.

### 2.3 Emergence of spontaneous physiological oscillations across frequency bands

Physiological oscillations across a range of frequency bands were observed in both the macaque and model thalamocortical circuits. In the model, these oscillations emerged despite having no oscillatory background inputs, suggesting they resulted from the intrinsic cellular biophysics and circuit connectivity. We quantified the power spectral density (PSD) of 10-second LFPs recorded from different macaques and from the model (Fig. 5). These results illustrate the high variability of spontaneous responses measured within and across macaques. This variability was comparable to that generated by the model. Despite the high variability, the model exhibited features similar to those observed consistently across macaques, including peaks at delta, theta/alpha and beta frequencies. To quantify these similarities and establish whether the LFP PSD generated by our model could be distinguished from that of macaques, we performed principal component (PCA) analysis (Fig. 5B). PCA explained a large proportion of the variance (PC1=57%, PC2=14%). As can be observed, the cluster of model data points partly (10/25 data points) overlapped those of macaques 1 and 2, yielding them indistinguishable via PCA (circled green points in Fig. 5B). Furthermore, the mean PCA distance between the macaque 2 and 3 clusters is greater (2.2) than the mean PCA distance between the model and macaque 2 clusters (1.3). Further validation was provided by plotting a shuffled version of the model LFP PSDs, which appeared as a clearly separate cluster with no overlap with the macaque data (with the exception of one outlier from macaque 3). The correlation matrix (Fig. 5C) across all LFP PSDs showed a much stronger correlation between the model and macaque than between shuffled model and macaque (0.31 vs 0.006; p<0.001, rank-sum test). Individual oscillation events were detected in current source density (CSD) data from resting state recordings gathered in silico from the A1 model, and in vivo from non-human primates (NHP), using software that has previously been used to detect and quantify features of oscillation events in human and NHP electrophysiology recordings (Neymotin, Barczak, et al. 2020). Once identified, oscillation events were classified according to frequency band: delta (0.5-4 Hz), theta (4-9 Hz), alpha (9-15 Hz), beta (15-29 Hz), gamma (30-80 Hz). Oscillation events were then sorted once more based on their laminar location, in either the supragranular, granular, or infragranular layers. We were thus able to compare model and NHP oscillation events that occurred in the same regions of the cortical column, within the same frequency band. Examples of matching individual oscillation events from each frequency band are shown in Fig. 6

**Figure 5.**
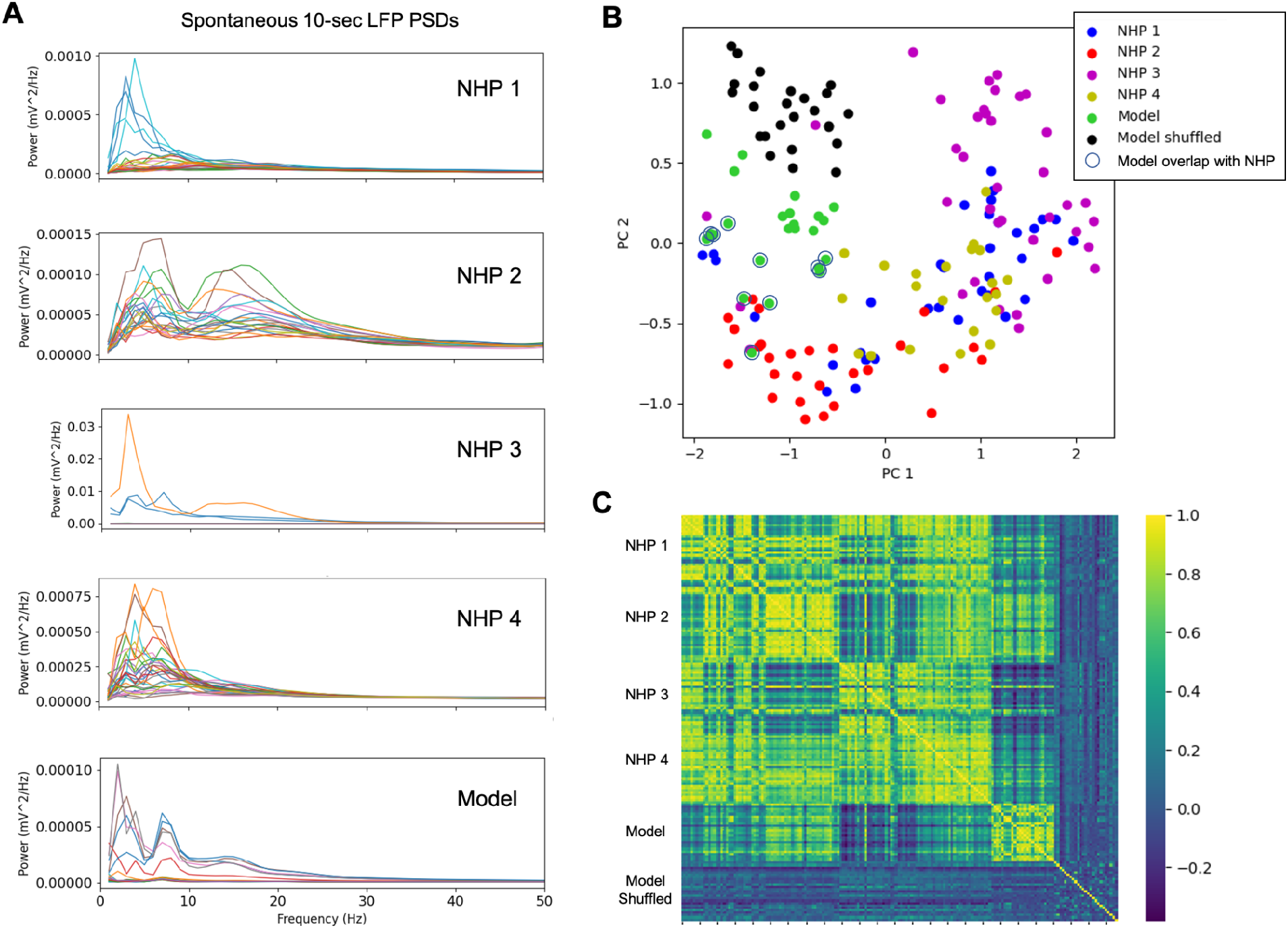
LFP power spectral density (PSD) of macaques and model. A) PSDs of 10-sec LFPs recorded from 4 macaques exhibit high variability within and across individuals; and show features consistent with the model LFP PSD, including peaks at delta, theta/alpha and beta. B) PCA analysis of the LFP PSDs reveal an overlap between model and macaque that is absent in the shuffled model. C) Correlation matrix of LFP PSDs illustrates the model is more strongly correlated with the macaques than the shuffled model.

**Figure 6.**
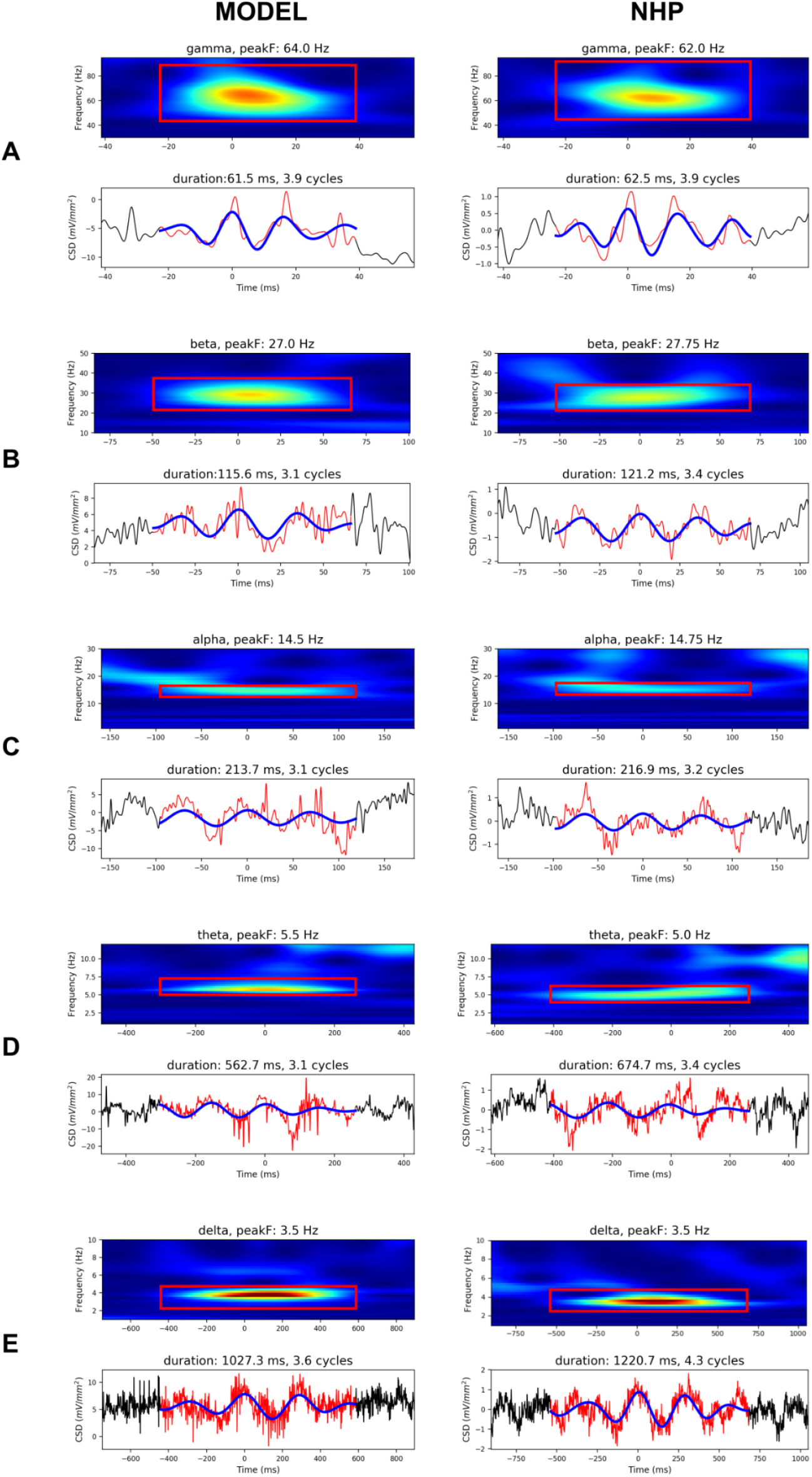
Similar oscillation events detected in CSD recorded from the model (left) and non-human primates (right), across different frequency bands. Each event is depicted with a spectrogram of the CSD data on the top panel (with a red bounding box delineating the oscillation event), and the CSD signal on the bottom panel (red: raw CSD time-series, blue: CSD time-series filtered with a bandpass filter with cutoffs at minimum, maximum event frequencies shown in spectrogram red bounding box). These examples demonstrate the model’s ability to reproduce physiologically realistic oscillation events across different frequency bands: A) Gamma B) Beta C) Alpha D) Theta E) Delta. (Note that the Theta and Alpha oscillations events are from supragranular layers, the Beta and Delta events are from infragranular layers, and the Gamma event is from the granular layer).

Several features were used to compare oscillation events across model and NHP data from different frequency bands, including temporal duration, peak frequency, and number of cycles in the oscillation (see Fig. 7). Overall, these three features showed similar average values and overlapping distributions when compared across the model and NHP, and across cortical layers. Duration was the most consistent value, with close average values across model and NHP at all frequency bands (p>0.05, t-test). Average peak frequency did not show significant differences for most frequency bands (p>0.05, t-test), with the exception of: 1) theta, which showed a slightly lower average frequency compared to the NHP data (p<0.05, t-test), and 2) gamma, which showed a slightly higher average frequency compared to NHP (p<0.05, t-test). Similarly, the number of cycles per oscillation event were on average the same across frequency bands (p>0.05, t-test), with the exception of gamma, which showed a slightly higher average value (p<0.05, t-test). The minor discrepancies in average values may however represent an artifact due the short overall duration of simulations (10 sec), compared to the duration of NHP recordings, which were on the order of minutes.

**Figure 7.**
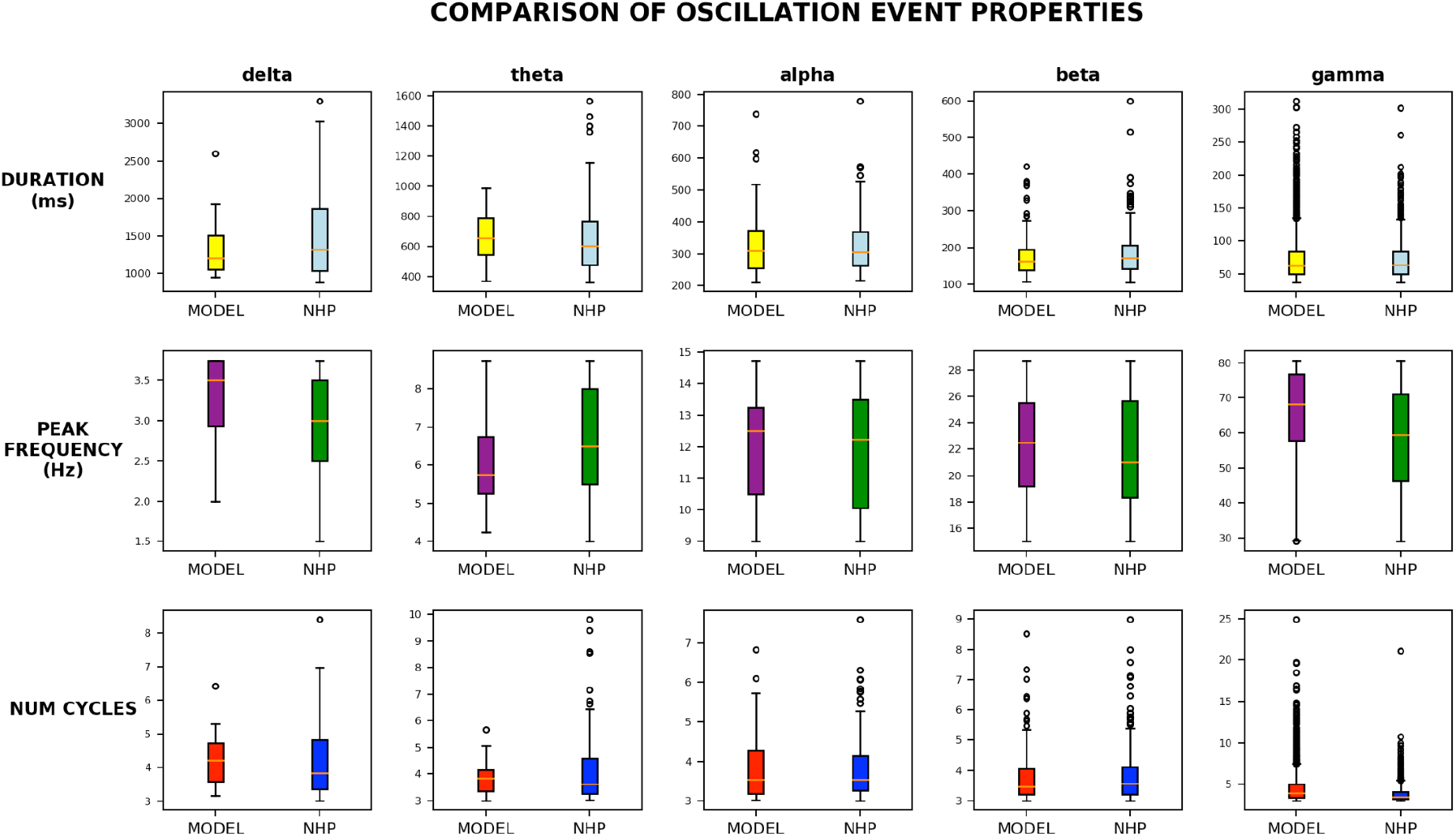
Oscillation event features in model and non-human primate data show strong similarity. Oscillation events were detected in resting state recordings, then sorted by frequency band (delta, theta, alpha, beta, gamma). The top row shows boxplots comparing the duration (ms) of the oscillation events in each frequency band, with model data shown in yellow, and NHP data in blue. Comparison of the peak frequency (Hz) of the model (purple) and NHP events (green) is shown in the middle row, while the number of cycles per event is compared in the bottom row, with model data in red and NHP data in blue. For each of the boxplots, the box itself ranges from the first to the third quartile, with the orange line depicting the median of the data set. The whiskers extend to 1.5 times the interquartile range, and outliers are represented with black circles. With the exception of minor differences in theta, gamma peak frequency, and gamma number of cycles, the average values of each feature were the same across model and NHP (t-test, p>0.05).

### 2.4 Unraveling the biophysical mechanisms underlying physiological oscillations at the cellular and circuit scales

After verifying that the oscillation events detected in the model data were comparable to the events observed in the NHP data, we used the model to examine the network- and population-level activity occurring during these oscillation events. This illustrates one of the advantages of the model. Not only were we able to generate the overall LFP and CSD data, but the biological detail of the model also allowed us to record and examine each population’s contribution to the overall LFP and CSD signals. Additionally, we were able to examine the spiking activity of each population at the time of each oscillation event, similar to multiunit activity observed during neurophysiological recordings, but with additional cell-type specificity.

Fig. 8 illustrates this approach using a physiologically realistic oscillation event detected in the simulated A1 column supragranular layer (Fig. 8A). As shown in Fig. 6, similar oscillation events were detected in the macaque A1 data. To determine the biophysical circuit sources underlying this oscillation event, we first examined the CSD signal generated by each cell population over all channels (Fig. 8B). This information showed that the layer 4 stellate (ITS4), layer 4 pyramidal tract (ITP4), and layer 5A intratelencephanic (IT5A) neural populations made the strongest contributions to the amplitude of the overall CSD signal during this event. Consistent with the model prediction, the dominant CSD peak frequencies of these three populations was very similar to that of the overall theta oscillation event (overall: 6.5 Hz (Fig. 8A); ITS4: 6.75 Hz (Fig. 8C), ITP4: 6.75 Hz (Fig. 8D); IT5A: 7 Hz (Fig. 8E). Interestingly, the individual population CSDs were not perfectly phase aligned; specifically, the IT5A signal (Fig. 8E) appeared to be shifted by approximately 10-20 ms with respect to the layer 4 populations (Fig. 8C,D). Although here we are only showing the top three contributing populations, other contributing populations also exhibited similar CSD signal phase shifts. We hypothesize that these phase shifts are responsible for the increased noise observed in the overall CSD signal compared to the individual population CSD signals that generated it. This increased noise may in turn explain the small differences in the CSD peak frequencies observed between the overall signal and the population signals that composed it. The contribution of IT5A to the overall CSD signal recorded at channel 8 is particularly interesting given that the IT5A cell somas are located at a cortical depth of 1250-1350 μm, whereas the channel 8 signals arise from electrodes at 700-900 μm depth. This suggests IT5A apical dendrite currents generate a substantial component of the detected CSD theta oscillation.

**Figure 8.**
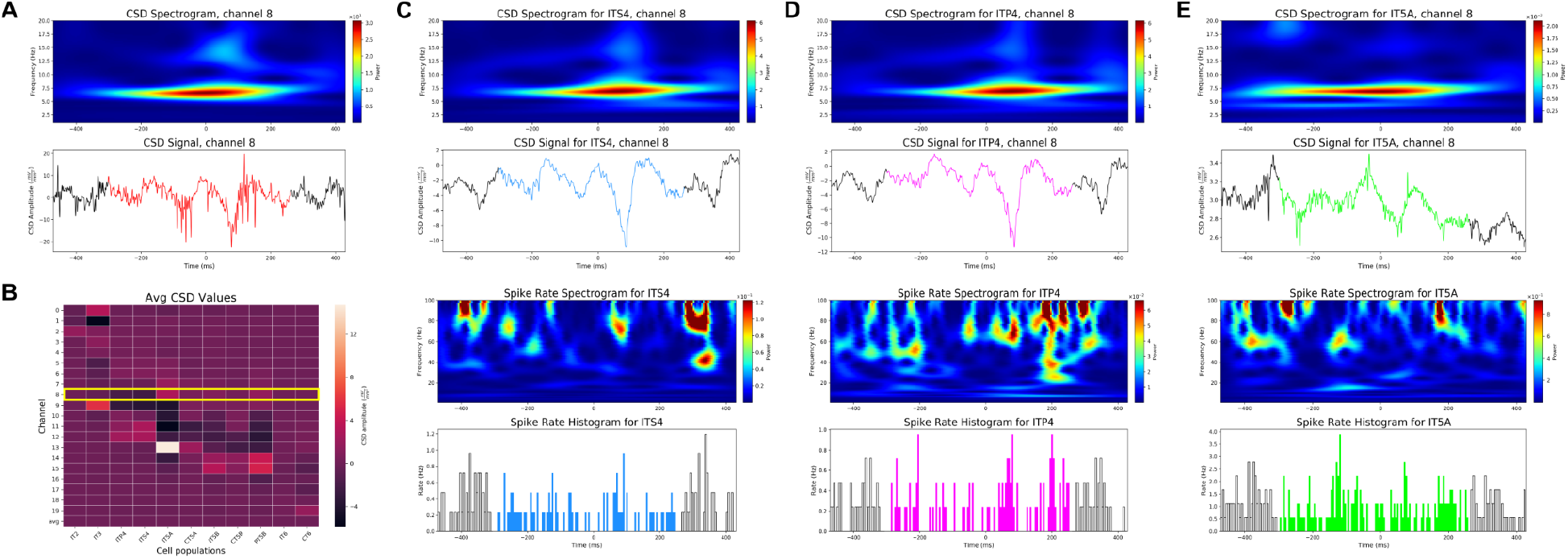
Model predicts layer- and cell type-specific sources of LFP/CSD during oscillations. **A)** CSD reveals a theta oscillation in A1 supragranular layer (see Fig. 6). **B)** Heat map depicting the average CSD amplitude (mV/mm^2^) during the time of the theta oscillation event across excitatory model populations and channels. The CSD spectrogram and time series (top panels) and the firing rate spectrogram and histogram (bottom panels) of the 3 populations with strongest contributions to the theta oscillation depicted in **(A)** are shown in: **(C)** ITS4, **(D)** ITP4, and **(E)** IT5A.

The corresponding spiking activity was then examined for each of these cell populations (Fig. 8C, D, E, bottom panels). Notably, the low-frequency theta event coincided with higher frequency gamma events in the spike rate spectrograms for all populations involved. These gamma events also occurred during periods when the population spike rates were elevated, consistent with the observation that elevated excitatory neuron firing leads to detectable gamma signals (Leszczyński et al. 2020). In addition, there was evidence of spike/field coherence, seen in the peak firing times of neural populations coinciding with peaks in the CSD theta rhythm. Overall, the presence of coincident theta and gamma demonstrates a cross-frequency interaction often observed in neural oscillations (Jensen and Colgin 2007), and highlights how the model can be used to make predictions on the origins of these complex dynamics observed in auditory cortex in vivo (O’Connell et al. 2015).

## 3. Discussion

### 3.1. Key findings and novelties

We have developed the first detailed multiscale model of macaque auditory thalamocortical circuits, including MGB, TRN and A1, and validated it against in vivo experimental data. The model integrated experimental data on the physiology, morphology, biophysics, density, laminar distribution and proportion of different cell types, as well as their local and long-range synaptic connectivity (Figs. 1-2). Realistic auditory inputs can be provided to the thalamus via a phenomenological model of the cochlear nucleus, auditory nerve and IC. The model generated cell type and layer-specific firing rates in the ranges observed experimentally, and accurately simulated the corresponding measures at multiple scales: local field potentials (LFPs), laminar current source density (CSD) analysis, and electroencephalogram (EEG) (Fig. 3). We identified multiple laminar CSD patterns during spontaneous activity and responses to speech similar to those recorded experimentally (Fig. 4). Physiological oscillations emerged across frequency bands without external rhythmic inputs and were comparable to those recorded spontaneously in vivo. Despite significant variability across animals and over time, the spectral power showed peaks at delta, theta/alpha and beta frequencies in all animals and in the model (Fig. 5). Additionally, individual CSD oscillation events closely matched physiological examples in all frequency bands (Fig. 6). Statistics on simulated CSD oscillation event duration, peak frequency and number of cycles were consistent across layers and frequency bands with those reported in vivo (Fig. 7). We used the model to make predictions about the contributions of distinct neural populations to specific oscillation events (Fig. 8). Notably, the model highlights the importance of disentangling individual neuronal population contributions to oscillatory activity. For example, the model predicted that the CSD theta oscillation event in Fig. 8A was caused predominantly by three layer 4 and 5 populations, each generating oscillatory activity with similar peak frequencies and small phase shifts. Taken together, our findings underline the significant role modeling plays when interpreting the basic properties of in vivo electrophysiology data.

Although circuit models of similar size and complexity have been developed, these models have largely focused on rodent visual (Billeh et al. 2020) and somatosensory (Markram et al. 2015) cortices. A highly-detailed rat somatosensory cortex model was used to study stimulus specific adaptation in the auditory cortex by modifying thalamic inputs (Amsalem et al. 2020), but the overall model cortical architecture and connectivity was not adapted to replicate the particularities of auditory circuits or the macaque species. Compared to our model, previous models of auditory cortex lack significant detail in terms of neuron model complexity, range of cell types, neuronal density and distribution, and/or circuit connectivity (Park and Geffen 2020; Stanley et al. 2019; Loebel, Nelken, and Tsodyks 2007; Zulfiqar, Moerel, and Formisano 2019; Kudela et al. 2018).

In short, the main novelties are that this model: 1) incorporates available data specific to the macaque species and auditory cortex; 2) includes a wide range of excitatory and inhibitory cell types from both cortical and thalamic regions; 3) uses synaptic connectivity that is cell type and layer-specific, and includes bidirectional thalamic connections with distinct core and matrix projections; 4) simulates realistic auditory inputs through a cochlear and inferior colliculus model; 5) generates realistic multiscale measures, including spiking activity, LFP, CSD, and EEG; and 6) recapitulates a range of macaque A1 in vivo results.

### 3.2. Challenges and limitations

Due to gaps in experimental data and in our theoretical understanding of biological principles, the model is necessarily incomplete and inaccurate and will need to be revised as more in vivo data becomes available. This is particularly true for the NHP auditory system, which has been less studied and is not as well characterized as, for example, the rodent visual system. Specifically, the availability of electrophysiological and connectivity data from the macaque auditory system for the different cortical and thalamic cell types was limited, so, when required, we used data from other macaque regions or from other mammalian auditory systems. Validating the layer and cell type-specific firing rates was also challenging due to lack of macaque A1 data; thus, many of the comparisons to experiments rely on the readily available laminar LFP and CSD measures. Despite these limitations, we believe our model incorporates more properties specific to macaque A1 and has been validated against more macaque A1 data than any previous model. Furthermore, it can be iteratively improved and further validated as newer and more precise data becomes available.

Generating physiologically constrained firing rates in all model populations required parameter tuning (also referred to as parameter fitting or optimization) of the connection strengths within biologically realistic ranges. When compared to our previous motor cortex model (Sivagnanam et al. 2020; Dura-Bernal et al. 2022), this process was particularly challenging in the NHP auditory system model, and required developing and iteratively improving our automated parameter optimization methods. We believe the reasons for this include the addition of two inhibitory cell types (VIP, NGF) and the incorporation of thalamic circuitry, which resulted in complex recurrent intracortical and thalamocortical interactions. The optimization methods resulted in a range of distinct model parameter combinations that produced valid network dynamics - a phenomenon known as parameter degeneracy. It is well known that biological neural circuits exhibit this same property: different combinations of neuron intrinsic and synaptic properties -- each varying up to several orders of magnitude -- can result in the circuit exhibiting the same physiological and functional outcome (Prinz, Bucher, and Marder 2004).

The relatively narrow diameter (200 um) of our simulated cortical column did not allow for a detailed implementation of the tonotopic organization of thalamic inputs. Nonetheless, the A1 column was tuned to a specific best frequency, as determined by the filtering of inputs through the cochlear and IC model. The A1 column also received a realistic number of afferent core and matrix thalamic inputs, with layer and cell type specificity. Future model versions can be extended to have a larger diameter column, or multiple columns, each receiving distinct thalamic projections, enabling the studying of circuit mechanisms that support frequency discrimination of auditory stimuli. Hence, in this study, we did not attempt to reproduce speech responses in detail, and instead focused on reproducing features of spontaneous activity, including the high variability observed experimentally.

We simulated, for the first time, EEG signals based on the current dipoles of individual neurons in a realistic model of macaque auditory cortical circuits. Calculating the voltage at the different scalp electrodes requires a realistic head volume conduction model. Unfortunately, we did not find a macaque head model, and had to use the standard human head model available within the LFPy tool (Hagen et al. 2018). This served as additional proof-of-concept of the multiscale capabilities of our model.

### 3.3. Outlook on research and clinical applications

Overall, the computational model provides a quantitative theoretical framework to integrate and interpret a wide range of experimental data, generate testable hypotheses and make quantitative predictions. It constitutes a powerful tool to study the biophysical underpinnings of different experimental measurements, including LFP, EEG, and MEG (Neymotin, Daniels, et al. 2020). This theoretical framework represents a baseline model that can be updated and extended as new data becomes available. Ongoing efforts by the BRAIN Initiative Cell Census Network (BICCN) and others may soon provide a cell census of the mammalian auditory cortex, similar to that recently made available for the motor cortex (BRAIN Initiative Cell Census Network (BICCN) 2021). Our model is fully open source and implemented using the NetPyNE tool (Dura-Bernal et al. 2019), which was explicitly designed to facilitate integration of experimental data through an intuitive language focused on describing biological parameters. This will enable other researchers to readily adapt the model to reproduce experimental manipulations, e.g. chemogenetic or pharmacologic interventions, or dynamics associated with different brain diseases. Work is already ongoing to adapt the model to study the EEG correlates of schizophrenia in A1 (Metzner and Steuber 2021), and to evaluate a novel LFP recording device (Abrego et al. 2021). To facilitate interoperability with other tools, NetPyNE can also export the model to standard formats, such as NeuroML (Gleeson et al. 2019) and SONATA (Dai et al. 2020), making it widely available to the community. This is also the first detailed circuit model to incorporate naturalistic auditory inputs, allowing future research linking structure, dynamics and function, and providing insights into neural representations during naturalistic stimulus processing. Given the general similarities between NHP and human thalamocortical circuitry (Herculano-Houzel 2009; Passingham 1973), this data-driven model has high translational relevance, and can start to bridge the gap across species and offer insights into healthy and pathological auditory system dynamics in humans.

## 4. Methods

We developed a model of the macaque auditory system consisting of a phenomenological model of cochlea and IC, and biophysically-detailed models of auditory thalamic and cortical circuits (Fig. 1). We validated the model against macaque in vivo experimental data. This section details the modeling, experimental and analysis methods used.

### 4.1 Single neuron models

#### Morphology and physiology of neuron classes

The network includes conductance-based cell models with parameters optimized to reproduce physiological responses. We used simplified morphologies of 1 - 6 compartments for each cell type, and sized dendritic lengths to match macaque cortical dimensions (Oliver et al. 2018, Figs. 8.3/4.4). We fitted the electrophysiological properties of each cell type to extant electrophysiology data from macaque when available, or other animal models when it was not. Passive parameters, such as membrane capacitance, were tuned to fit resting membrane potential (RMP) and other features of subthreshold traces (e.g. sag from hyperpolarization). Active parameters included values such as the fast sodium channel density, and were tuned to reproduce characteristics like oscillatory bursting and firing rate vs input current (f-I) curve (see Supplementary Fig. 1).

Within the A1 network, we modeled four classes of excitatory neurons: the intratelencephalic spiny stellate (ITS), intratelencephalic pyramidal (IT), pyramidal tract (PT) and corticothalamic (CT). These were distributed across the six cortical layers. The ITS model consisted of 3 compartments (a soma and 2 dendrites), and was adapted from a previously published Layer 4 spiny stellate model (Mainen and Sejnowski 1996). There is evidence for the presence of stellate cells in A1 in mammals, including rodents, rabbits, bats, cats and humans (Meyer, González-Hernández, and Ferres-Torres 1989; Oliver et al. 2018; Harris and Shepherd 2015; Y. Wang, Brzozowska-Prechtl, and Karten 2010), although in some species these were relatively rare compared to visual and somatosensory cortices. Several macaque studies also mention the role of A1 L4 stellate cells in receiving input from thalamus (Steinschneider et al. 1998, 1992; Fishman et al. 2000). The IT, PT, and CT cell models were each composed of 6 compartments: a soma, axon, basal dendrite, and 3 apical dendrites. These models were based on previous work (Neymotin et al. 2017), in which simplified cell models were optimized to reproduce subthreshold and firing dynamics observed in vivo (Suter, Migliore, and Shepherd 2013; Yamawaki et al. 2014; Oswald et al. 2013). Apical dendrite lengths were modified to match macaque cortical dimensions and layer-specific connectivity requirements. The classification of cortical neurons into IT, PT and CT was based not only on their projection targets, but also on their local connectivity, laminar location, morphology, intrinsic physiology and genetics (Harris and Shepherd 2015; Shepherd and Yamawaki 2021). Although the PT terminology may be confusing for A1, this cell class refers to subcerebral projection neurons, including brainstem, and has been previously used for non-motor cortical regions (A1, V1, S1) (Harris and Shepherd 2015; Shepherd and Yamawaki 2021; Baker et al. 2018). PT neurons have also been labeled as ‘extratelencephalic’ (ET), but this does not distinguish them from the also extratelencephalic CT neurons. In A1, a category of neurons described as ‘large pyramidal cells’ overlap significantly with features of the PT cell class: mostly occupy L5B, have thick-tufted morphologies reaching up to L1, are intrinsically bursting and project to brainstem, including inferior colliculus (IC), superior olivary complex and the cochlear nuclear complex (Budinger and Kanold 2018; Jeffery A. Winer et al. 2002; Baker et al. 2018; Jeffery A. Winer and Schreiner 2010).

Four classes of inhibitory neurons (NGF, SOM, PV, VIP) were also simulated in the A1 network model. The vasoactive intestinal peptide (VIP) cell model was based on a previously published 5-compartment model (Turi et al. 2019), whereas the somatostatin (SOM) and parvalbumin (PV) interneurons were based on published 3-compartment models (Konstantoudaki et al. 2014). Parameters such as dendritic length were modified to better fit extant cortical data regarding rheobase and f-I curve (Tripathy et al. 2015). The neurogliaform (NGF) cell model was adapted from an existing model in rodent (Bezaire et al. 2016), with soma compartment size modified to more closely match the geometry of NGF cells in monkeys (Povysheva et al. 2007). Channel mechanisms, including A-type potassium and Ih currents, were also added to the soma compartment to replicate the electrophysiological characteristics (e.g. sag, f-I curve) described for these cell types in the literature (Povysheva et al. 2007).

In thalamus, the modeled MGB contained thalamocortical (TC) cells, high-threshold thalamocortical cells (HTC), and local thalamic interneurons (TI). The TC and HTC cells were both single-compartment models capable of tonic and burst firing (Iavarone et al. 2019), with the HTC model having the addition of a high-threshold T-type channel mechanism (Vijayan and Kopell 2012). The locally inhibitory TI cells had 2 compartments (a soma and a dendrite) and were fitted to in vitro electrophysiology data recorded from lateral geniculate nucleus (Zhu, Uhlrich, and Lytton 1999a, [b] 1999; Zhu, Lytton, and Xue 1999). These cells were optimized to reproduce the oscillatory bursting observed in this cell type (Zhu, Lytton, and Xue 1999). The thalamic reticular nucleus (TRN) contained the single-compartment inhibitory reticular (IRE) cells, with parameters also optimized to display this cell type’s characteristic intrinsic rhythmicity (Destexhe et al. 1994, 1996).

### 4.2 Thalamocortical circuit model populations

#### Auditory thalamus

Our auditory thalamus model included the medial geniculate body (MGB) and the thalamic reticular nucleus (TRN). The MGB was composed of two types of thalamocortical neurons (TC, HTC) and thalamic interneurons (TI). TRN was composed of reticular nucleus cells (IRE). The overall proportion of excitatory to inhibitory neurons was 3:1. For TC, TI and IRE cell types, we included two separate populations in order to capture the distinct connectivity patterns of the core vs matrix thalamic circuits. Matrix populations were labeled with an ‘M’ at the end: TCM, IREM, TIM. The proportion of core to matrix neurons was 1:1 (Bonjean et al. 2012). The density and ratio of the different thalamic populations was based on experimental data (J. A. Winer and Larue 1996; Huang, Larue, and Winer 1999). The resulting ratio of thalamic to cortical neurons was 1:17, consistent with published data (Coen-Cagli, Kanitscheider, and Pouget 2017).

#### Auditory cortex

We modeled a cylindrical volume of the macaque primary auditory cortex (A1) with a 200 µm diameter and 2000 µm height (cortical depth) including 12,187 neurons and over 25 million synapses (Fig. 1B). The cylinder diameter was chosen to approximately match the horizontal dendritic span of a neuron located at the center, consistent with previous modeling approaches (Markram et al. 2015; Billeh et al. 2020). Macaque cortical depth and layer boundaries were based on macaque published data (Kelly and Hawken 2017; Tremblay, Lee, and Rudy 2016). The model includes 36 neural populations distributed across the 6 cortical layers and consisting of 4 excitatory (IT, ITS, PT, CT), and 4 inhibitory types (SOM, PV, VIP and NGF). Details of the biophysics and morphology of each cell type are provided in section [“Single Neuron Models”] above. The laminar distribution, cell density and proportion of each cell type was based on experimental data (Kelly and Hawken 2017; Lefort et al. 2009; Schuman et al. 2019; Harris and Shepherd 2015). Layer 1 included only NGF cells. Layers 2 to 6 included IT, SOM, PV, VIP and NGF cells. Additionally, ITS cells were added to layer 4, PT cells to layer 5B, and CT cells to layers 5A, 5B and 6. The resulting number of cells in each population depended on the modeled volume, layer boundaries and neuronal proportions and densities per layer.

### 4.3 Thalamocortical circuit model connectivity

#### Connectivity parameters: connection probability and weight

We characterized connectivity in the thalamocortical circuit using two parameters for each projection: probability of connection and unitary connection strength. Probability of connection was defined as the probability that each neuron in the postsynaptic population was connected to a neuron in the presynaptic population. For example, if both pre- and postsynaptic populations have 100 neurons, a probability of 10% will result in an average of 1,000 connections (10% of the total 10,000 possible connections). The set of presynaptic neurons to connect to was randomly selected and autapses and multapses were not allowed. Given the neuronal morphologies were simplified to 6 or less compartments, we used a single synaptic contact for each cell-to-cell connection.

Unitary connection strength was defined as the EPSP amplitude in response to a spike from a single presynaptic neuron. Given that synaptic weights in NEURON are typically defined as a change in conductance (in uS), we derived a scaling factor to map unitary EPSP amplitude (in mV) to synaptic weights. To do this, we simulated an excitatory synaptic input to generate a somatic EPSP of 0.5 mV at each neuron segment. We then calculated a scaling factor for each neuron segment that converted the EPSP amplitude (mV) values used to define connectivity in NetPyNE into the corresponding NEURON synaptic weights (in uS). This resulted in the somatic EPSP response to a unitary connection input being independent of synaptic location, also termed synaptic democracy (Poirazi and Papoutsi 2020). This is consistent with experimental evidence showing synaptic conductances increased with distance from soma, “counterbalancing the filtering effects of the dendrites and reducing the location dependence of somatic EPSP amplitude” (Magee and Cook 2000). We thresholded dendritic scaling factors to 4x that of the soma to avoid overexcitability in the network in cases when neurons receive hundreds of inputs that interact nonlinearly (Spruston 2008; Behabadi et al. 2012).

#### Types of synapses

Excitatory synapses consisted of colocalized AMPA (rise, decay τ: 0.05, 5.3 ms) and NMDA (rise, decay τ: 15, 150 ms) receptors, both with reversal potentials of 0 mV. The ratio of NMDA to AMPA receptors was 1:1 (Myme et al. 2003), meaning their weights were each set to 50% of the connection weight. NMDA conductance was scaled by 1/(1+0.28 · Mg · e^(−0062 · V)^) with Mg = 1mM (Jahr and Stevens 1990). Inhibitory synapses from SOM to excitatory neurons consisted of a slow GABA_A_ receptor (rise, decay τ: 2, 100 ms) and GABA_B_ receptor, with a 9:1 ratio. Synapses from SOM to inhibitory neurons only included the slow GABA_A_ receptor. Synapses from PV consisted of a fast GABA_A_ receptor (rise, decay τ: 0.07, 18.2). Synapses from VIP included a different fast GABA_A_ receptor (rise, decay τ: 0.3, 6.4) (Pi et al. 2013), and synapses from NGF included the GABA_A_ and GABA_B_ receptors with a 1:1 ratio. The reversal potential was 0 mV from AMPA and NMDA, -80 mV for all GABA_A_ and -93 mV for GABA_B_. The GABA_B_ synapse was modeled using second messenger connectivity to a G protein-coupled, inwardly-rectifying potassium channel (GIRK) (Destexhe, Babloyantz, and Sejnowski 1993). The remaining synapses were modeled with a double-exponential mechanism.

#### Connection delays

Connection delays were estimated as 2 ms to account for presynaptic release and postsynaptic receptor delays, plus a variable propagation delay calculated as the 3D Euclidean distance between the pre- and postsynaptic cell bodies divided by a propagation speed of 0.5 m/s. Conduction velocities of unmyelinated axons range between 0.5-10 m/s (Purves et al. 2018), but here we chose the lowest value given that our soma-to-soma distance underestimates the non-straight trajectory of axons and the distance to target dendritic synapses.

#### Intra-thalamic connectivity

Intrathalamic connectivity was derived from existing rodent, cat and primate experimental and computational studies (Bonjean et al. 2012; Cruikshank et al. 2010; Serkov and Gonchar 1996; Jones 2002; Billeh et al. 2020) (see Fig. 2). More specifically, connection probabilities and unitary strength for TC→RE, RE→TC and RE→RE (both core and matrix populations) were largely based on a previous primate thalamus study (Bonjean et al. 2012) and validated with data from mouse ventrobasal thalamus (Cruikshank et al. 2010) and cat MGBv (Bonjean et al. 2012; Cruikshank et al. 2010; Serkov and Gonchar 1996; Jones 2002; Billeh et al. 2020). No evidence was found for TC recurrent connections. Thalamic interneuron connectivity was derived from the same cat MGBv study, which provided the number of synaptic contacts for TI→TI, TI→TC and TC→TI, from which we estimated the probability of connection from each projection. We also verified that our model intra-thalamic connectivity was generally consistent with that of the Allen Brain Institute visual thalamocortical model (Jones 2002; Billeh et al. 2020). Given that thalamic neuron models were single-compartment, no specific dendritic synaptic location information was included.

#### Intra-cortical connectivity

Connectivity within the A1 local circuit populations was defined as a function of pre- and postsynaptic cell type and layer. Given the overall lack of detailed cell type-specific connectivity experimental data for macaque A1, we used as a starting point the connectivity from two experimentally-grounded mammalian cortical microcircuit modeling studies: the Allen Brain Institute (ABI) V1 (Billeh et al. 2020) and the Blue Brain Project (BBP) S1 (Markram et al. 2015). We then updated the model connectivity with experimental data specific to macaque A1, when available, or simply mammalian A1.

Both studies included the projection-specific probability of connection and unitary connection strength parameters that we required for our model. However, the ABI V1 model had fewer excitatory (1) and inhibitory (3) broad cell types than our A1 model (4 E and 4 I), whereas the BBP S1 model included significantly more (11 E and 15 I). Neither model included the distinction between L5A and L5B present in our A1 model. The ABI V1 did provide length constants to implement distance-dependent connectivity, which we wanted to include for some of the A1 projections. Therefore, as a first step, we mapped our cell types to the closest ones in the ABI V1 model and obtained the corresponding connectivity matrices for A1. We then updated the A1 connectivity of cell types that were missing from ABI V1 based on data from BBP S1, more specifically, the ITS, PT, CT and VIP cell types. To do this we mapped A1 cell types to those closest in BBP S1, and scaled the connectivity parameters of missing cell types proportionally, using shared cell types as reference (e.g. IT or PV). Through this systematic approach we were able to combine data from ABI V1 and BBP S1 in a consistent way, to determine the connectivity parameters of all the A1 populations.

Inhibitory connections were further refined using data from A1 (Budinger and Kanold 2018; Kato, Asinof, and Isaacson 2017; Pi et al. 2013) or from studies with more detailed cell type-specific data (Tremblay, Lee, and Rudy 2016; Naka and Adesnik 2016). We updated the L2/3 SOM connectivity so they projected strongly not only to superficial layer excitatory neurons, but also to deeper ones by targeting their apical dendrites; this was not the case for PV cells, which projected strongly mostly to intralaminar excitatory neurons (Kato, Asinof, and Isaacson 2017; Naka and Adesnik 2016). More specifically, probabilities of connection from L2/3 SOM and PV to excitatory neurons were a function of the postsynaptic neuron layer (L1-L6) based on data from an A1 study (Kato, Asinof, and Isaacson 2017). The probability of connection from VIP to excitatory neurons was set to a very low value derived from mouse A1 data (Pi et al. 2013). Following this same study, VIP→SOM connections were made strong, VIP→PV weak, and VIP→VIP very weak. Connection probabilities of all I→E/I projections decayed exponentially with distance using a projection-specific length constant obtained from the ABI V1 study.

Information on the dendritic location of synaptic inputs was also incorporated, when available, into the model. Cortical excitatory synapses targeted the soma and proximal dendrites of L2-4 excitatory neurons, distal dendrites of L5-6 excitatory neurons, and were uniformly distributed in cortical inhibitory neurons (Billeh et al. 2020; Budinger and Kanold 2018; Harris and Shepherd 2015). L1 NGF neurons targeted the apical tuft of excitatory neurons, L2-4 NGF targeted the apical trunk of L2-4 excitatory neurons and the upper trunk of L5-6 excitatory neurons, and L5-6 NGF targeted the lower trunk of L5-6 excitatory neurons (Tremblay, Lee, and Rudy 2016; Budinger and Kanold 2018; Naka and Adesnik 2016). Synapses from SOM interneurons were uniformly distributed along excitatory neurons, those from PV and VIP neurons targeted the soma and proximal dendrites of excitatory neurons (Naka and Adesnik 2016; Kato, Asinof, and Isaacson 2017; Tremblay, Lee, and Rudy 2016).

#### Thalamocortical and corticothalamic connectivity

Thalamocortical connections were layer- and cell type-specific and were derived from studies in mouse auditory cortex (Ji et al. 2016) and rodent somatosensory cortex (Constantinople and Bruno 2013; Cruikshank et al. 2010). Core MGB thalamocortical neurons projected to cortical excitatory neurons in cortical layers 3 to 6. The strongest projections were to layer 4 ITP, ITS and PV neurons. Weaker thalamocortical projections also targeted L3 IT and PV; L4 SOM and NGF; L5-6 IT, CT and PV; and L5B PT and SOM. Matrix thalamocortical neurons projected to excitatory neurons in all layers except 4, and to L1 NGF, L2/3 PV and SOM, and L5-6 PV. Core thalamic inputs targeted the soma and proximal dendrites of cortical excitatory cells, whereas matrix thalamic inputs targeted their distal dendrites (Bonjean et al. 2012; Jones 2002).

Corticothalamic projections originating from L5A, L5B and L6 CT neurons targeted all core thalamus populations (TC, HTC, TI and IRE); whereas projections from L5B IT and PT neurons targeted the matrix thalamus populations (TCM, TIM, IREM). Connectivity data was derived from primate and rodent studies on auditory cortex and other cortical regions (Bonjean et al. 2012; Yamawaki and Shepherd 2015; Budinger and Kanold 2018; Harris and Shepherd 2015; Jones 2002; Crandall, Cruikshank, and Connors 2015).

### 4.4 Background inputs

To model the influence of the other brain regions not explicitly modeled on auditory cortex and thalamus, we provided background inputs to all our model neurons. These inputs were modeled as independent Poisson spike generators for each cell, targeting apical excitatory and basal inhibitory synapses, with an average firing rate of 40 Hz. Connection weights were automatically adjusted for each cell type to ensure that, in the absence of local circuit connectivity, all neurons exhibited a low spontaneous firing rate of approximately 1 Hz.

### 4.5 Full model synaptic weight tuning

#### Overview of approach

Although we followed a systematic data-driven approach to build our model, the complete experimental dataset required to build a detailed model of the macaque auditory thalamocortical system is currently not available. Therefore, we had to combine experimental data from different species, different brain regions and obtained using different recording techniques. It is therefore not surprising that in order to obtain physiologically constrained firing rates across all populations, we needed to tune the connectivity parameters. Automated optimization methods have been previously used for simpler networks (e.g. recurrent point-neuron spiking networks) (Nicola and Clopath 2017; Sussillo and Abbott 2009; Dura-Bernal et al. 2017; Carlson et al. 2014; Hasegan et al. 2021). However, optimization of large-scale biophysically-detailed networks typically requires expert-guided parameter adjustments (Bezaire et al. 2016; Markram et al. 2015), for example through parameter sweeps (grid search) (Billeh et al. 2020). In order to find a more systematic approach to tune this type of model, here we explored automated optimization methods, and gradually refined them and combined them with heuristic approaches as needed. Here we describe the final approach employed to obtain the tuned network.

#### Automated optimization algorithm

Our starting point was the network with cell type-specific background inputs, so that all cells fired at approximately 1 Hz in the absence of connectivity. We then added connectivity with parameters taken from the literature and similar existing models. The resulting network included many silent populations (0 Hz) and others firing at very high rates (>100 Hz). Our aim was to obtain a baseline network where all populations fired within biologically constrained rates.

After classical grid search methods failed, we evaluated the Optuna (http://optuna.org) (Akiba et al. 2019), a hyperparameter optimization framework designed for machine learning applications, which dynamically searches the parameter space. Compared to evolutionary algorithms we used in the past, Optuna has the advantage of producing similar results while not requiring all candidates of a generation to be completed before moving to the next one. Instead, it dynamically decides the next candidate to explore based on all the evaluated candidates up to that point, which makes it faster and less resource-consuming.

In order to automate the process, we implemented a fitness function to automatically evaluate how good each of the solutions was:

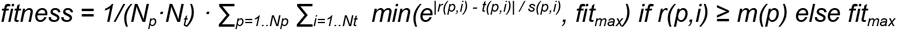

where *N*_*p*_ is the number of neural populations, *N*_*t*_ is the number of time periods that are evaluated, *p* is the population index, *i* is the time period index, *r(p, i)* is the average firing rate for population with index *p* during time period with index *i, t(p, i) and s(p, i)* are the target rate mean and standard deviation for population *p* and time period with index *i*, and *fit*_*max*_ is the maximum (worst possible) fitness value. For each population the target firing rate is described by a Gaussian function with mean and standard deviation and a minimum threshold (for E pops: mean=5, std=20, min=0.05; for I pops: mean=10, std=30, min=0.05). This Gaussian function is evaluated across four consecutive 250ms periods, to ensure relative homogeneity in the firing rates, e.g. to avoid populations firing only during the first 100ms, but with an average firing rate matching the target rate. The mean output of this function across populations and time periods constitutes the fitness error (Supplementary Fig. 2).

#### Layer-specific and cell-type specific parameters

In order to reduce the fitness errors, we gradually included more tuned parameters (see Fig. 11A). Our final approach included 4 projection-class weight gains (E→E, E→I, I→E, I→I) for each of the 7 layers (1, 2, 3, 4, 5A, 5B, 6). Analysis also revealed the highly specific dynamics for each of the four inhibitory cell types, which prompted us to include inhibitory cell type-specific weight gains: E→ PV, E→ SOM, E→ VIP, E→ NGF and PV→ E, SOM→ E, VIP→ E, NGF→ E. Including both layer-specific and cell type-specific parameters resulted in overall better solutions with lower fitness errors.

#### Stepwise layer-by-layer tuning

Increasing the number of parameters (dimensions) increases the size of the parameter space to explore, which increases the number of optimization trials (simulations) required to obtain a good solution, and increases the risk of getting stuck in local minima. There are two main ways to reduce the parameter space: 1) reducing the number of parameters, e.g. including only parameters for a subset of layers, or of projection types; and 2) reducing the range of parameter values explored, e.g. constraining these based on previous optimization results. Both of these solutions are implemented in the stepwise layer-by-layer tuning approach we employed, which reduced the massive HPC resources required to explore the large model parameter spaces.

To implement the layer-by-layer tuning approach, we first optimized the parameters within L4 alone. Once this layer achieved valid firing rates in all cell populations we added L3, and tuned the L3 connectivity parameters, while we kept L2 parameters within a small range of the previously obtained solution. We repeated this for L2, L5A, L5B, L6 and finally L1. Due to a small bug when tuning L2 and L3, once the full model was tuned, we retuned to L2 and L3 while keeping the rest of parameters within a small range (Fig. 11). A similar layer-by-layer approach was followed to tune the Allen Brain Institute V1 model (Billeh et al. 2020), although they used a heuristic unidimensional grid search approach, whereas we employed an automated multidimensional dynamic search using Optuna.

#### Projection-specific weight tuning

Once we had obtained a reasonable solution for most model populations using the layer-by-layer approach, additional fine-tuning was required to improve the rate of specific populations. In particular, the SOM2 and SOM3 were 0 Hz and PV2 and VIP2 were firing too high (>100 Hz). The current parameters explored did not appear to provide enough specificity to improve the rate of these populations without worsening some of the others. Therefore, we had to tune the weight gains of specific population-to-population projections, e.g. from IT2 to SOM2. Using Optuna, we optimized the weights of all projections targeting the populations with inadequate rates: PV2, SOM2, VIP2 and SOM3. This resulted in improved rates for these populations.

#### Final model

Our final network included all 43 thalamic and cortical populations firing within 0.1 and 25 Hz, i.e. no epileptic or silent populations. Due to the unprecedented scale and level of detail in the model, e.g. complex interaction between 4 interneuron types, we had to employ an exploratory approach evaluating many several methods to tune the weights. Overall, this required over 500,000 simulations and over 5 million core hours on Google Cloud HPCs. The lessons learned during this process should facilitate the automated tuning of similar detailed models in the future.

### 4.6 Phenomenological models of peripheral auditory structures

To simulate spontaneous activity in our baseline model we used background white noise as inputs to our thalamic and cortical populations. However, in order to accurately simulate auditory stimuli input we also connected a model of peripheral auditory structures such as the auditory nerve (AN) and inferior colliculus (IC). To simulate these structures, we used phenomenological models that captured the signal transformations occurring in these regions (Krishna and Semple 2000). These models produced outputs that were used to drive the thalamocortical cells in the downstream, more biologically detailed portion of the auditory pathway model. The AN responses modeled here included several characteristic nonlinearities such as rate saturation, adaptation, and phase locking (Carney, Li, and McDonough 2015; Krishna and Semple 2000). Outputs from the AN model were convolved and modulated with synaptic information and used as inputs to a phenomenological model of inferior colliculus (IC). Model neurons of the IC utilized different types of modulation transfer functions to capture both the spectral and amplitude modulation tuning observed in this structure (Carney, Li, and McDonough 2015; Nelson and Carney 2004; Krishna and Semple 2000; Joris, Schreiner, and Rees 2004). These phenomenological models mitigated common encoding issues encountered at high frequencies and high sound levels, providing us with IC outputs that were useful throughout a broad range of frequencies and noise (Carney, Li, and McDonough 2015).

The AN and IC models were implemented in Matlab and are available within the UR_EAR 2.0 tool (see Supplementary Fig. 3). We used .wav files as input to this tool and obtained time-resolved IC firing rates. The model allowed customization of several options, including the cochlear central frequency and bandwidth. We saved the firing rates for different input sounds and converted these to spike times using a Python-based inhomogeneous Poisson generator (Muller et al. 2007). We then used NEURON spike generators (VecStims) defined in NetPyNE to provide the IC spike times as input to the model thalamic populations.

### 4.7 Model building, simulation and optimization

We developed the computational model using the NetPyNE tool (Dura-Bernal et al. 2019), and ran all parallel simulations using NEURON 8.0 (Lytton et al. 2016; Carnevale and Hines 2006) with a fixed time step of 0.05 ms. NetPyNE is a python package that provides a high-level interface to NEURON, and allows for the definition of complicated multiscale models using an intuitive declarative language focused on the biological parameters. NetPyNE then translates these specifications into a NEURON model, facilitates running parallel simulations, and automates the optimization and exploration of parameters using supercomputers. We executed our simulations primarily on Google Cloud supercomputers using a Slurm-based cluster with 80-core compute nodes (Sivagnanam et al. 2020). Some simulations were also run on XSEDE supercomputers Comet and Stampede, either using our own allocations or through the Neuroscience Gateway (NSG) (Sivagnanam et al. 2013). We used the NetPyNE software tool to design, execute, organize, and analyze the simulations, as well as to export our model to the SONATA (Dai et al. 2020) and NeuroML (Gleeson et al. 2019) standards.

### 4.8 Data Analysis and Visualization

#### Spiking raster plot, firing rate statistics and voltage traces

The NetPyNE package (Dura-Bernal et al. 2019) was used to record and analyze simulation output data, and to visualize spiking raster plots, firing rate statistics, and neuronal membrane voltage traces.

#### Local Field Potential (LFP)

At each in silico electrode, the local field potential (LFP) was calculated as the sum of the extracellular potential from each neuronal segment. For estimation of extracellular potential, we used the line source approximation method and assumed that the model neurons were immersed in an ohmic medium with a fixed conductivity of sigma = 0.3 mS/mm (Parasuram et al. 2016; Gold et al. 2006; Dura-Bernal et al. 2019). Electrodes were spatially distributed at 100 μm intervals along a vertical axis of the 2000 μm A1 column. Model LFP recording, analysis and visualization was performed using the NetPyNE package.

#### Current Source Density (CSD)

We compared the in silico current source density (CSD) signals with in vivo data recorded from the supragranular, granular, and infragranular layers of A1 while NHPs were at rest. CSD was calculated as the second spatial derivative of the LFP. CSD analysis and visualization was performed using the NetPyNE package (Dura-Bernal et al. 2019).

#### Oscillation event detection

Using the OEvent package (Neymotin, Barczak, et al. 2020), we also used Morlet wavelet spectrograms and their corresponding CSD waveforms to identify individual oscillation events occurring in spontaneous data, and to compare these events across in vivo and in silico contexts. OEvent extracted moderate/high-power events using 7-cycle Morlet wavelets on non-overlapping 10 s windows (Sherman et al. 2016; Neymotin, Daniels, et al. 2020). We used linearly spaced frequencies (0.25 Hz frequency increments) ranging from 0.25 - 125 Hz. Power time-series of each wavelet transform were normalized by median power from the recording/simulation. We applied a local maximum filter to detect peaks in the spectrogram. Local peaks were assessed to determine whether their power exceeded a 4x median threshold to detect moderate- to high-power events. Frequency and time bounds around the peak were determined by including time and frequency values before/after, above/below peak frequency until power fell below the smaller of ½ maximum event amplitude and 4× median threshold. As shown in Fig. 6, this produced a bounding box around each oscillation event that was used to determine frequency spread (minF to maxF), duration, and peak frequency (frequency at which maximum power is detected). We merged events when their bounding box overlapping area in the spectrogram exceeded 50% of the minimum area of each individual event. This allowed for the continuity of events separated by minor fluctuations below threshold. We then calculated additional features from this set of events, including the number of cycles (event duration × peak frequency). We classified events into standard frequency bands on the following intervals: delta (0.5-4 Hz), theta (4-9 Hz), alpha (9-15 Hz), beta (15-29 Hz), gamma (30-80 Hz). Classification was based on the frequency at which maximum power occurred during each event. These oscillation event analysis techniques yielded morphologically similar events between the simulated and NHP data (Figs. 6, 7). Non-normalized CSD data were used to validate and analyze the contributions of individual cell populations to the detected oscillation events (Fig. 8).

### 4.9 Experiment recordings

We used a dataset which included local field potentials invasively recorded from the primary auditory cortex (A1) of 4 female non-human primates as they sat quietly in a dark room with their eyes mostly open (previously described in (Neymotin, Barczak, et al. 2020)). In a subset of recordings, short sentences in the English language were presented at 80dB SPL. In both conditions, there were no behavioral requirements of the NHPs and no rewards were offered. Outside of the recording sessions, NHPs had full access to fluids and food.

All procedures were approved in advance by the Animal Care and Use Committee of the Nathan Kline Institute. NHP data was recorded during acute penetrations of A1 in rhesus macaques weighing 5-8 kg, who had been prepared surgically for chronic awake electrophysiological recordings. Prior to surgery, each animal was adapted to a custom fitted primate chair and to the sound proofed recording chamber. Surgical preparation was performed under general anesthesia using aseptic techniques (for details see (Schroeder 1998; Lakatos et al. 2013)). Briefly, to provide access to the brain, either Cilux (Crist Instruments) or Polyetheretherketone (PEEK; Rogue Research Inc.) recording chambers were positioned normal to the cortical surface of the superior temporal plane for orthogonal penetration of A1. These recording chambers and a PEEK headpost (used to permit painless head restraint) were secured to the skull with ceramic screws and embedded in dental acrylic. Each NHP was given a minimum of 6 weeks for post-operative recovery before behavioral training and data collection began.

During recordings, NHPs were head-fixed and linear array multielectrodes (23 contacts with 100, 125 or 150µm intercontact spacing, Plexon Inc.) were acutely positioned to sample all cortical layers of A1. Neuroelectric signals were continuously recorded with a sampling rate of 44 kHz using the Alpha Omega SnR system. For NHP data analyses using current-source density (CSD) signals, CSD was calculated as the second spatial derivative of laminar local field potential. This was done to reduce potential issues related to volume conducted activity.

## Acknowledgements

Research supported by NIDCD R01DC012947, NIBIB U24EB028998, NIMH P50MH109429, NIBIB U01EB017695, NYS DOH01-C32250GG-3450000, NSF 190444, Army Research Office W911NF-19-1-0402, ARO URAP supplement. The views and conclusions contained in this document are those of the authors and should not be interpreted as representing the official policies, either expressed or implied, of the Army Research Office or the U.S. Government. The U.S. Government is authorized to reproduce and distribute reprints for Government purposes, notwithstanding any copyright notation herein.

## Supplementary Figures

**Supplementary Figure 1.**
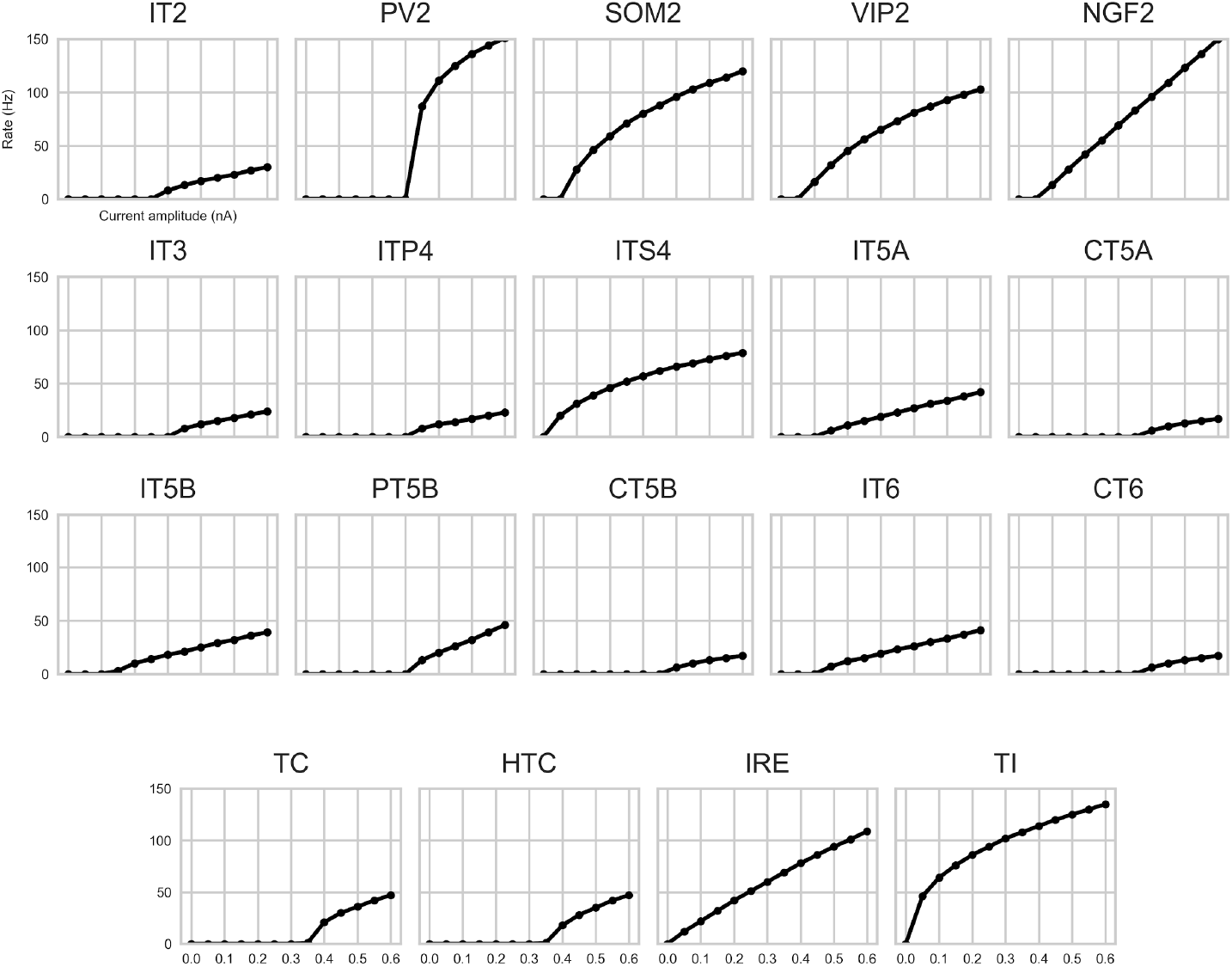
Frequency-current (f-I) curves of all distinct cortical and thalamic cell types used in the model. X-axis shows the amplitude of somatic current injection provided over a 1 second interval, and y-axis shows the number of action potentials produced.

**Supplementary Figure 2.**
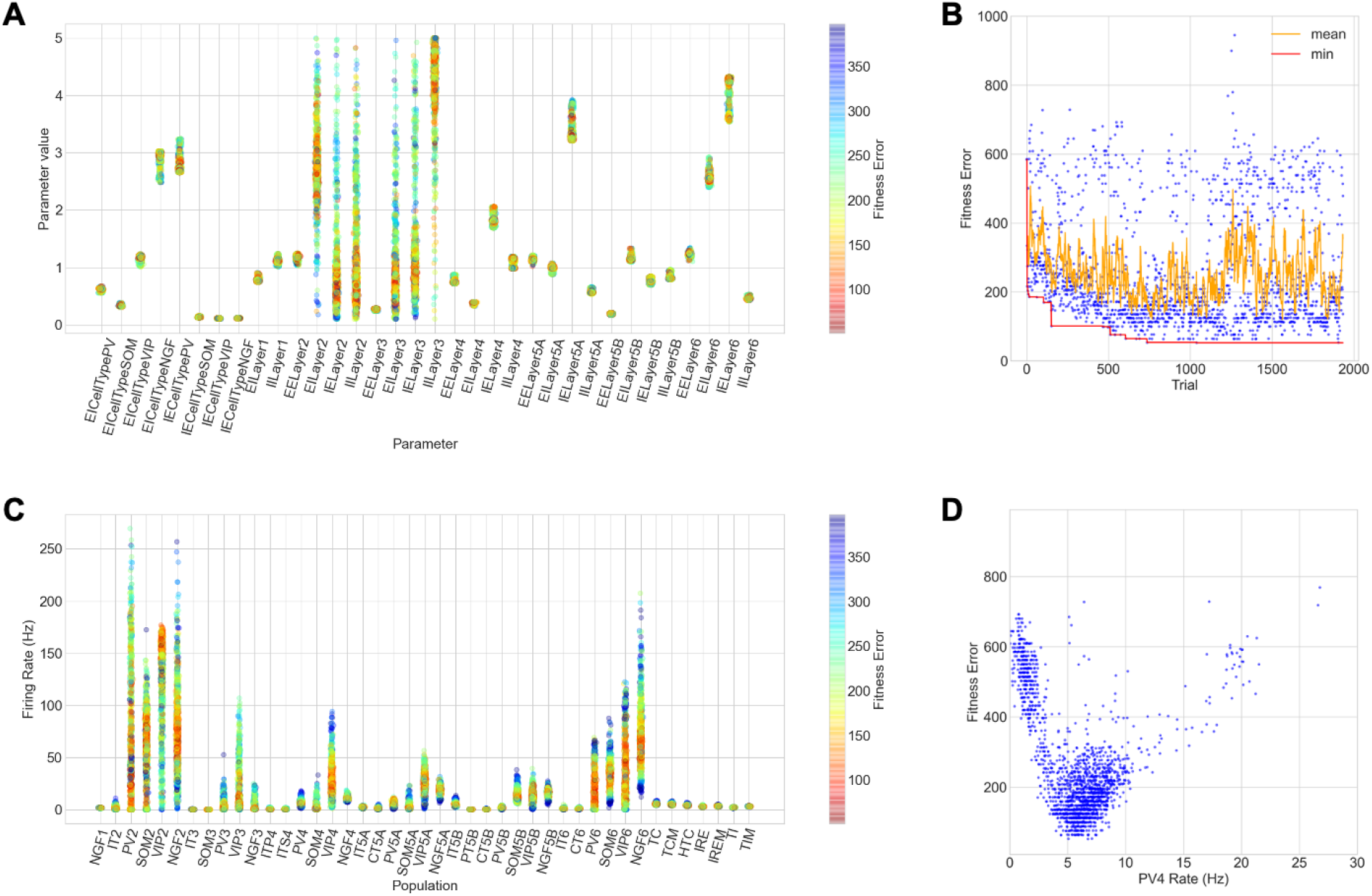
Optimization of connectivity parameters in auditory thalamocortical model. A) Range of values explored for each parameter tuned, and fitness error (color) associated with each value. Only parameter values with fitness errors below 400 were included; red indicates approximate final parameter value. Note that only layer 2 and 3 connectivity parameters were tuned across a wide value range, while the rest of parameters were highly constrained based on previous optimizations. B) Fitness error of the trials, each evaluating a different parameter combination; moving mean average across 10 trials (orange); overall minimum fitness (red) shows fast improvement up to ∼750 trials and then plateaus. C) Range of average firing rates obtained for each population, and fitness error (color) associated with each value. Only rates with fitness errors below 400 were included; red indicates approximate final rate. D) Relation between fitness error and PV4 average firing rate; the V-shape indicates very low or very high firing rates were correlated with high fitness errors, whereas firing rates close to the target (∼5-10 Hz) were correlated with low fitness error.

**Supplementary Figure 3.**
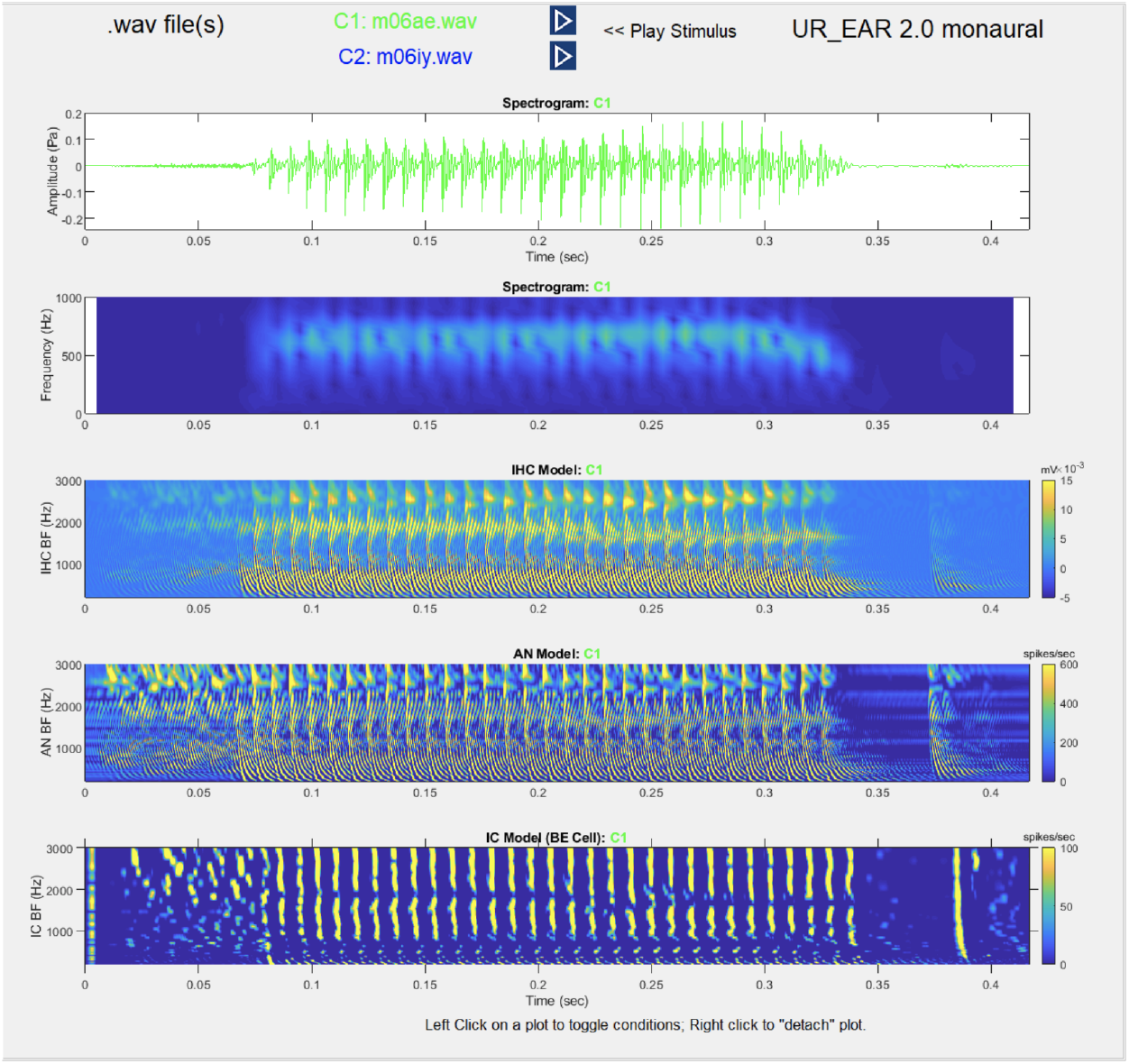
Example output of the cochlea and inferior colliculus (IC) phenomenological model. From top to bottom: time series of example input sound (.wav file); spectrogram of the input sound; output of the Cochlear inner hair cells (IHC) model (in mV); output of the auditory nerve (AN) model (in Hz); output of the IC model (in mV). The firing rates of the IC units are then converted to spike times and provided as input to the biophysically detailed MGB model.

## References

Abrego, Amada M., Wasif Khan, Christopher E. Wright, M. Rabiul Islam, Mohammad H. Ghajar, Xiaokang Bai, Nitin Tandon, and John P. Seymour. 2021. “Sensing Local Field Potentials with a Directional and Scalable Depth Array: The DISC Electrode Array.” bioRxiv. https://doi.org/10.1101/2021.09.20.460996.

Ahveninen, Jyrki, Norbert Kopčo, and Iiro P. Jääskeläinen. 2014. “Psychophysics and Neuronal Bases of Sound Localization in Humans.” Hearing Research 307 (January): 86–97.

Akiba, Takuya, Shotaro Sano, Toshihiko Yanase, Takeru Ohta, and Masanori Koyama. 2019. “Optuna: A Next-Generation Hyperparameter Optimization Framework.” In Proceedings of the 25th ACM SIGKDD International Conference on Knowledge Discovery & Data Mining, 2623–31. KDD ‘19. New York, NY, USA: Association for Computing Machinery.

Albouy, Philippe, Jérémie Mattout, Romain Bouet, Emmanuel Maby, Gaëtan Sanchez, Pierre-Emmanuel Aguera, Sébastien Daligault, et al. 2013. “Impaired Pitch Perception and Memory in Congenital Amusia: The Deficit Starts in the Auditory Cortex.” Brain: A Journal of Neurology 136 (Pt 5): 1639–61.

Amsalem, Oren, James King, Michael Reimann, Srikanth Ramaswamy, Eilif Muller, Henry Markram, Israel Nelken, and Idan Segev. 2020. “Dense Computer Replica of Cortical Microcircuits Unravels Cellular Underpinnings of Auditory Surprise Response.” BioRxiv. https://doi.org/10.1101/2020.05.31.126466.

Andéol, Guillaume, Anne Guillaume, Christophe Micheyl, Sophie Savel, Lionel Pellieux, and Annie Moulin. 2011. “Auditory Efferents Facilitate Sound Localization in Noise in Humans.” The Journal of Neuroscience: The Official Journal of the Society for Neuroscience 31 (18): 6759–63.

Baguley, David M. 2003. “Hyperacusis.” Journal of the Royal Society of Medicine 96 (12): 582–85.

Baker, Arielle, Brian Kalmbach, Mieko Morishima, Juhyun Kim, Ashley Juavinett, Nuo Li, and Nikolai Dembrow. 2018. “Specialized Subpopulations of Deep-Layer Pyramidal Neurons in the Neocortex: Bridging Cellular Properties to Functional Consequences.” The Journal of Neuroscience: The Official Journal of the Society for Neuroscience 38 (24): 5441–55.

Behabadi, Bardia F., Alon Polsky, Monika Jadi, Jackie Schiller, and Bartlett W. Mel. 2012. “Location-Dependent Excitatory Synaptic Interactions in Pyramidal Neuron Dendrites.” PLoS Computational Biology 8 (7): e1002599.

Bezaire, Marianne J., Ivan Raikov, Kelly Burk, Dhrumil Vyas, and Ivan Soltesz. 2016. “Interneuronal Mechanisms of Hippocampal Theta Oscillation in a Full-Scale Model of the Rodent CA1 Circuit.” eLife 5: e18566.

Billeh, Yazan N., Binghuang Cai, Sergey L. Gratiy, Kael Dai, Ramakrishnan Iyer, Nathan W. Gouwens, Reza Abbasi-Asl, et al. 2020. “Systematic Integration of Structural and Functional Data into Multi-Scale Models of Mouse Primary Visual Cortex.” Neuron 106 (3): 388–403.e18.

Bonjean, Maxime, Tanya Baker, Maxim Bazhenov, Sydney Cash, Eric Halgren, and Terrence Sejnowski. 2012. “Interactions between Core and Matrix Thalamocortical Projections in Human Sleep Spindle Synchronization.” The Journal of Neuroscience: The Official Journal of the Society for Neuroscience 32 (15): 5250–63.

BRAIN Initiative Cell Census Network (BICCN). 2021. “A Multimodal Cell Census and Atlas of the Mammalian Primary Motor Cortex.” Nature 598 (7879): 86–102.

Brody, Robert M., Brian D. Nicholas, Michael J. Wolf, Paula B. Marcinkevich, and Gregory J. Artz. 2013. “Cortical Deafness: A Case Report and Review of the Literature.” Otology & Neurotology: Official Publication of the American Otological Society, American Neurotology Society [and] European Academy of Otology and Neurotology 34 (7): 1226–29.

Budinger, Eike, and Patrick O. Kanold. 2018. “Auditory Cortex Circuits.” In The Mammalian Auditory Pathways: Synaptic Organization and Microcircuits, edited by Douglas L. Oliver, Nell B. Cant, Richard R. Fay, and Arthur N. Popper, 199–233. Cham: Springer International Publishing.

Carlile, Simon, Russell Martin, and Ken McAnally. 2005. “Spectral Information in Sound Localization.” In International Review of Neurobiology, 70:399–434. Academic Press.

Carlson, Kristofor David, Jayram M. Nageswaran, Nikil Dutt, and Jeffrey L. Krichmar. 2014. “An Efficient Automated Parameter Tuning Framework for Spiking Neural Networks.” Frontiers in Neuroscience 8 (10). https://doi.org/10.3389/fnins.2014.00010.

Carnevale, Nicholas T., and Michael L. Hines. 2006. The NEURON Book. Cambridge University Press.

Carney, Laurel H., Tianhao Li, and Joyce M. McDonough. 2015. “Speech Coding in the Brain: Representation of Vowel Formants by Midbrain Neurons Tuned to Sound Fluctuations.” eNeuro 2 (4). https://doi.org/10.1523/ENEURO.0004-15.2015.

Cavinato, Marianna, Jessica Rigon, Chiara Volpato, Carlo Semenza, and Francesco Piccione. 2012. “Preservation of Auditory P300-like Potentials in Cortical Deafness.” PloS One 7 (1): e29909.

Coen-Cagli, Ruben, Ingmar Kanitscheider, and Alexandre Pouget. 2017. “A Method to Estimate the Number of Neurons Supporting Visual Orientation Discrimination in Primates.” F1000Research 6 (September): 1752.

Constantinople, Christine M., and Randy M. Bruno. 2013. “Deep Cortical Layers Are Activated Directly by Thalamus.” Science 340 (6140): 1591–94.

Crandall, Shane R., Scott J. Cruikshank, and Barry W. Connors. 2015. “A Corticothalamic Switch: Controlling the Thalamus with Dynamic Synapses.” Neuron 86 (3): 768–82.

Cruikshank, Scott J., Hayato Urabe, Arto V. Nurmikko, and Barry W. Connors. 2010. “Pathway-Specific Feedforward Circuits between Thalamus and Neocortex Revealed by Selective Optical Stimulation of Axons.” Neuron 65 (2): 230–45.

Dai, Kael, Juan Hernando, Yazan N. Billeh, Sergey L. Gratiy, Judit Planas, Andrew P. Davison, Salvador Dura-Bernal, et al. 2020. “The SONATA Data Format for Efficient Description of Large-Scale Network Models.” PLoS Computational Biology 16 (2): e1007696.

De Stefano, Lisa A., Lauren M. Schmitt, Stormi P. White, Matthew W. Mosconi, John A. Sweeney, and Lauren E. Ethridge. 2019. “Developmental Effects on Auditory Neural Oscillatory Synchronization Abnormalities in Autism Spectrum Disorder.” Frontiers in Integrative Neuroscience 13 (July): 34.

Destexhe, A., A. Babloyantz, and T. J. Sejnowski. 1993. “Ionic Mechanisms for Intrinsic Slow Oscillations in Thalamic Relay Neurons.” Biophysical Journal 65: 1538–52.

Destexhe, A., T. Bal, D. A. McCormick, and T. J. Sejnowski. 1996. “Ionic Mechanisms Underlying Synchronized Oscillations and Propagating Waves in a Model of Ferret Thalamic Slices.” Journal of Neurophysiology 76 (3): 2049–70.

Destexhe, A., D. Contreras, T. J. Sejnowski, and M. Steriade. 1994. “A Model of Spindle Rhythmicity in the Isolated Thalamic Reticular Nucleus.” Journal of Neurophysiology 72 (2): 803–18.

Dimitrijevic, Andrew, Michael L. Smith, Darren S. Kadis, and David R. Moore. 2017. “Cortical Alpha Oscillations Predict Speech Intelligibility.” Frontiers in Human Neuroscience 11 (February): 88.

Dura-Bernal, Salvador, Samuel A. Neymotin, C. C. Kerr, Subhashini Sivagnanam, A. Majumdar, J. T. Francis, and William W. Lytton. 2017. “Evolutionary Algorithm Optimization of Biological Learning Parameters in a Biomimetic Neuroprosthesis.” IBM Journal of Research and Development 61 (2/3): 6:1–6:14.

Dura-Bernal, Salvador, Samuel A. Neymotin, Benjamin A. Suter, Joshua Dacre, Julia Schiemann, Ian Duguid, Gordon M. G. Shepherd, and William W. Lytton. 2022. “Multiscale Model of Primary Motor Cortex Circuits Reproduces in Vivo Cell Type-Specific Dynamics Associated with Behavior.” bioRxiv. https://doi.org/10.1101/2022.02.03.479040.

Dura-Bernal, Salvador, Benjamin A. Suter, Padraig Gleeson, Matteo Cantarelli, Adrian Quintana, Facundo Rodriguez, David J. Kedziora, et al. 2019. “NetPyNE, a Tool for Data-Driven Multiscale Modeling of Brain Circuits.” eLife 8 (April). https://doi.org/10.7554/eLife.44494.

Dykstra, Andrew R., Christine K. Koh, Louis D. Braida, and Mark Jude Tramo. 2012. “Dissociation of Detection and Discrimination of Pure Tones Following Bilateral Lesions of Auditory Cortex.” PloS One 7 (9): e44602.

Edgar, J. Christopher, Sarah Y. Khan, Lisa Blaskey, Vivian Y. Chow, Michael Rey, William Gaetz, Katelyn M. Cannon, et al. 2015. “Neuromagnetic Oscillations Predict Evoked-Response Latency Delays and Core Language Deficits in Autism Spectrum Disorders.” Journal of Autism and Developmental Disorders 45 (2): 395–405.

Eggermont, J. J. 1992. “Stimulus Induced and Spontaneous Rhythmic Firing of Single Units in Cat Primary Auditory Cortex.” Hearing Research 61 (1-2): 1–11.

Fishman, Y. I., D. H. Reser, J. C. Arezzo, and M. Steinschneider. 2000. “Complex Tone Processing in Primary Auditory Cortex of the Awake Monkey. I. Neural Ensemble Correlates of Roughness.” The Journal of the Acoustical Society of America 108 (1): 235–46.

Fontolan, L., B. Morillon, C. Liegeois-Chauvel, and Anne-Lise Giraud. 2014. “The Contribution of Frequency-Specific Activity to Hierarchical Information Processing in the Human Auditory Cortex.” Nature Communications 5 (September): 4694.

Gandal, Michael J., J. Christopher Edgar, Richard S. Ehrlichman, Mili Mehta, Timothy P. L. Roberts, and Steven J. Siegel. 2010. “Validating γ Oscillations and Delayed Auditory Responses as Translational Biomarkers of Autism.” Biological Psychiatry 68 (12): 1100–1106.

Ghitza, Oded. 2011. “Linking Speech Perception and Neurophysiology: Speech Decoding Guided by Cascaded Oscillators Locked to the Input Rhythm.” Frontiers in Psychology 2 (June): 130.

Giraud, Anne-Lise, and David Poeppel. 2012. “Cortical Oscillations and Speech Processing: Emerging Computational Principles and Operations.” Nature Neuroscience 15 (4): 511–17.

Gleeson, Padraig, Matteo Cantarelli, Boris Marin, Adrian Quintana, Matt Earnshaw, Sadra Sadeh, Eugenio Piasini, et al. 2019. “Open Source Brain: A Collaborative Resource for Visualizing, Analyzing, Simulating, and Developing Standardized Models of Neurons and Circuits.” Neuron, June. https://doi.org/10.1016/j.neuron.2019.05.019.

Gold, C., D. A. Henze, C. Koch, and G. Buzsaki. 2006. “On the Origin of the Extracellular Action Potential Waveform: A Modeling Study.” Jnphys 95 (5): 3113–28.

Gourévitch, Boris, Régine Le Bouquin Jeannès, Gérard Faucon, and Catherine Liégeois-Chauvel. 2008. “Temporal Envelope Processing in the Human Auditory Cortex: Response and Interconnections of Auditory Cortical Areas.” Hearing Research 237 (1-2): 1–18.

Hagen, Espen, Solveig Næss, Torbjørn V. Ness, and Gaute T. Einevoll. 2018. “Multimodal Modeling of Neural Network Activity: Computing LFP, ECoG, EEG, and MEG Signals With LFPy 2.0.” Frontiers in Neuroinformatics 12 (December): 92.

Hamilton, Liberty S., Yulia Oganian, Jeffery Hall, and Edward F. Chang. 2021. “Parallel and Distributed Encoding of Speech across Human Auditory Cortex.” Cell 184 (18): 4626–39.e13.

Harris, Kenneth D., and Gordon M. G. Shepherd. 2015. “The Neocortical Circuit: Themes and Variations.” Nature Neuroscience 18 (2): 170–81.

Hasegan, Daniel, Matt Deible, Christopher Earl, David D’Onofrio, Hananel Hazan, Haroon Anwar, and Samuel A. Neymotin. 2021. “Multi-Timescale Biological Learning Algorithms Train Spiking Neuronal Network Motor Control.” bioRxiv. https://doi.org/10.1101/2021.11.20.469405.

Herculano-Houzel, Suzana. 2009. “The Human Brain in Numbers: A Linearly Scaled-up Primate Brain.” Frontiers in Human Neuroscience 3 (November): 31.

Hirano, Shogo, Alexander Nakhnikian, Yoji Hirano, Naoya Oribe, Shigenobu Kanba, Toshiaki Onitsuka, Margaret Levin, and Kevin M. Spencer. 2018. “Phase-Amplitude Coupling of the Electroencephalogram in the Auditory Cortex in Schizophrenia.” Biological Psychiatry. Cognitive Neuroscience and Neuroimaging 3 (1): 69–76.

Hirano, Yoji, Naoya Oribe, Toshiaki Onitsuka, Shigenobu Kanba, Paul G. Nestor, Taiga Hosokawa, Margaret Levin, Martha E. Shenton, Robert W. McCarley, and Kevin M. Spencer. 2020. “Auditory Cortex Volume and Gamma Oscillation Abnormalities in Schizophrenia.” Clinical EEG and Neuroscience: Official Journal of the EEG and Clinical Neuroscience Society 51 (4): 244–51.

Holmes, Emma, and Ingrid S. Johnsrude. 2021. “Speech-Evoked Brain Activity Is More Robust to Competing Speech When It Is Spoken by Someone Familiar.” NeuroImage 237 (August): 118107.

Hromádka, Tomás, Michael R. Deweese, and Anthony M. Zador. 2008. “Sparse Representation of Sounds in the Unanesthetized Auditory Cortex.” PLoS Biology 6 (1): e16.

Huang, C. L., D. T. Larue, and J. A. Winer. 1999. “GABAergic Organization of the Cat Medial Geniculate Body.” The Journal of Comparative Neurology 415 (3): 368–92.

Hyde, Krista L., Isabelle Peretz, and Robert J. Zatorre. 2008. “Evidence for the Role of the Right Auditory Cortex in Fine Pitch Resolution.” Neuropsychologia 46 (2): 632–39.

Iavarone, Elisabetta, Jane Yi, Ying Shi, Bas-Jan Zandt, Christian O’Reilly, Werner Van Geit, Christian Rössert, Henry Markram, and Sean L. Hill. 2019. “Experimentally-Constrained Biophysical Models of Tonic and Burst Firing Modes in Thalamocortical Neurons.” PLoS Computational Biology 15 (5): e1006753.

Jahr, C. E., and C. F. Stevens. 1990. “Voltage Dependence of NMDA-Activated Macroscopic Conductances Predicted by Single-Channel Kinetics.” Jnsci 10 (9): 3178–82.

Jensen, Ole, and Laura L. Colgin. 2007. “Cross-Frequency Coupling between Neuronal Oscillations.” Trends in Cognitive Sciences 11 (7): 267–69.

Ji, Xu-Ying, Brian Zingg, Lukas Mesik, Zhongju Xiao, Li I. Zhang, and Huizhong W. Tao. 2016. “Thalamocortical Innervation Pattern in Mouse Auditory and Visual Cortex: Laminar and Cell-Type Specificity.” Cerebral Cortex 26 (6): 2612–25.

Jochaut, Delphine, Katia Lehongre, Ana Saitovitch, Anne-Dominique Devauchelle, Itsaso Olasagasti, Nadia Chabane, Monica Zilbovicius, and Anne-Lise Giraud. 2015. “Atypical Coordination of Cortical Oscillations in Response to Speech in Autism.” Frontiers in Human Neuroscience 9 (March): 171.

Jones, Edward G. 2002. “Thalamic Circuitry and Thalamocortical Synchrony.” Philosophical Transactions of the Royal Society of London. Series B, Biological Sciences 357 (1428): 1659–73.

Joris, P. X., C. E. Schreiner, and A. Rees. 2004. “Neural Processing of Amplitude-Modulated Sounds.” Physiological Reviews 84 (2): 541–77.

Kato, Hiroyuki K., Samuel K. Asinof, and Jeffry S. Isaacson. 2017. “Network-Level Control of Frequency Tuning in Auditory Cortex.” Neuron 95 (2): 412–23.e4.

Kelly, Jenna G., and Michael J. Hawken. 2017. “Quantification of Neuronal Density across Cortical Depth Using Automated 3D Analysis of Confocal Image Stacks.” Brain Structure & Function 222 (7): 3333–53.

Konstantoudaki, Xanthippi, Athanasia Papoutsi, Kleanthi Chalkiadaki, Panayiota Poirazi, and Kyriaki Sidiropoulou. 2014. “Modulatory Effects of Inhibition on Persistent Activity in a Cortical Microcircuit Model.” Frontiers in Neural Circuits 8 (January): 7.

Krishna, B. S., and M. N. Semple. 2000. “Auditory Temporal Processing: Responses to Sinusoidally Amplitude-Modulated Tones in the Inferior Colliculus.” Journal of Neurophysiology 84 (1): 255–73.

Kudela, Pawel, Dana Boatman-Reich, David Beeman, and William Stanley Anderson. 2018. “Modeling Neural Adaptation in Auditory Cortex.” Frontiers in Neural Circuits 12 (September): 72.

Lakatos, Peter, Annamaria Barczak, Samuel A. Neymotin, Tammy McGinnis, Deborah Ross, Daniel C. Javitt, and Monica Noelle O’Connell. 2016. “Global Dynamics of Selective Attention and Its Lapses in Primary Auditory Cortex.” Nature Neuroscience, September. https://doi.org/10.1038/nn.4386.

Lakatos, Peter, Gabriella Musacchia, Monica Noelle O’Connell, Arnaud Y. Falchier, Daniel C. Javitt, and Charles E. Schroeder. 2013. “The Spectrotemporal Filter Mechanism of Auditory Selective Attention.” Neuron 77 (4): 750–61.

Lakatos, Peter, Monica Noelle O’Connell, Annamaria Barczak, Tammy McGinnis, Samuel A. Neymotin, Charles E. Schroeder, John F. Smiley, and Daniel C. Javitt. 2019. “The Thalamocortical Circuit of Auditory Mismatch Negativity.” Biological Psychiatry 0 (0). https://doi.org/10.1016/j.biopsych.2019.10.029.

Latinus, Marianne, Phil McAleer, Patricia E. G. Bestelmeyer, and Pascal Belin. 2013. “Norm-Based Coding of Voice Identity in Human Auditory Cortex.” Current Biology: CB 23 (12): 1075–80.

Lefort, Sandrine, Christian Tomm, J-C Floyd Sarria, and Carl C. H. Petersen. 2009. “The Excitatory Neuronal Network of the C2 Barrel Column in Mouse Primary Somatosensory Cortex.” Neuron 61 (2): 301–16.

Leszczyński, Marcin, Annamaria Barczak, Yoshinao Kajikawa, Istvan Ulbert, Arnaud Y. Falchier, Idan Tal, Saskia Haegens, Lucia Melloni, Robert T. Knight, and Charles E. Schroeder. 2020. “Dissociation of Broadband High-Frequency Activity and Neuronal Firing in the Neocortex.” Science Advances 6 (33): eabb0977.

Loebel, Alex, Israel Nelken, and Misha Tsodyks. 2007. “Processing of Sounds by Population Spikes in a Model of Primary Auditory Cortex.” Frontiers in Neuroscience 1 (1): 197–209.

Lytton, William W., Alexandra Seidenstein, Salvador Dura-Bernal, Felix Schurmann, Robert A. McDougal, and Michael L. Hines. 2016. “Simulation Neurotechnologies for Advancing Brain Research: Parallelizing Large Networks in NEURON.” Neural Computation 28: 2063–90.

Magee, Jeffrey C., and Erik P. Cook. 2000. “Somatic EPSP Amplitude Is Independent of Synapse Location in Hippocampal Pyramidal Neurons.” Nature Neuroscience 3 (9): 895–903.

Mainen, Z. F., and T. J. Sejnowski. 1996. “Influence of Dendritic Structure on Firing Pattern in Model Neocortical Neurons.” Nature 382 (6589): 363–66.

Markov, N. T., P. Misery, A. Falchier, C. Lamy, J. Vezoli, R. Quilodran, M. A. Gariel, et al. 2011. “Weight Consistency Specifies Regularities of Macaque Cortical Networks.” Cerebral Cortex 21 (6): 1254–72.

Markram, Henry, Eilif Muller, Srikanth Ramaswamy, Michael W. Reimann, Marwan Abdellah, Carlos Aguado Sanchez, Anastasia Ailamaki, et al. 2015. “Reconstruction and Simulation of Neocortical Microcircuitry.” Cell 163 (2): 456–92.

Matsumoto, Riki, Hisaji Imamura, Morito Inouchi, Tomokazu Nakagawa, Yohei Yokoyama, Masao Matsuhashi, Nobuhiro Mikuni, et al. 2011. “Left Anterior Temporal Cortex Actively Engages in Speech Perception: A Direct Cortical Stimulation Study.” Neuropsychologia 49 (5): 1350–54.

Merchant, Saumil N., and John J. Rosowski. 2008. “Conductive Hearing Loss Caused by Third-Window Lesions of the Inner Ear.” Otology & Neurotology: Official Publication of the American Otological Society, American Neurotology Society [and] European Academy of Otology and Neurotology 29 (3): 282–89.

Metzner, Christoph, and Volker Steuber. 2021. “The Beta Component of Gamma-Band Auditory Steady-State Responses in Patients with Schizophrenia.” Scientific Reports 11 (1): 20387.

Meyer, G., T. H. González-Hernández, and R. Ferres-Torres. 1989. “The Spiny Stellate Neurons in Layer IV of the Human Auditory Cortex. A Golgi Study.” Neuroscience 33 (3): 489–98.

Muller, Eilif, Lars Buesing, Johannes Schemmel, and Karlheinz Meier. 2007. “Spike-Frequency Adapting Neural Ensembles: Beyond Mean Adaptation and Renewal Theories.” Neural Computation 19 (11): 2958–3010.

Myme, Chaelon I. O., Ken Sugino, Gina G. Turrigiano, and Sacha B. Nelson. 2003. “The NMDA-to-AMPA Ratio at Synapses Onto Layer 2/3 Pyramidal Neurons Is Conserved Across Prefrontal and Visual Cortices.” Journal of Neurophysiology 90 (2): 771–79.

Naka, Alexander, and Hillel Adesnik. 2016. “Inhibitory Circuits in Cortical Layer 5.” Frontiers in Neural Circuits 10 (35). https://doi.org/10.3389/fncir.2016.00035.

Nelson, Paul C., and Laurel H. Carney. 2004. “A Phenomenological Model of Peripheral and Central Neural Responses to Amplitude-Modulated Tones.” The Journal of the Acoustical Society of America 116 (4 Pt 1): 2173–86.

Neymotin, Samuel A., Annamaria Barczak, Monica Noelle O’Connell, Tammy McGinnis, Noah Markowitz, Elizabeth Espinal, Erica Y. Griffith, et al. 2020. “Taxonomy of Neural Oscillation Events in Primate Auditory Cortex.” bioRxiv. https://www.biorxiv.org/content/10.1101/2020.04.16.045021v1.abstract.

Neymotin, Samuel A., Dylan S. Daniels, Blake Caldwell, Robert A. McDougal, Nicholas T. Carnevale, Mainak Jas, Christopher I. Moore, Michael L. Hines, Matti Hämäläinen, and Stephanie R. Jones. 2020. “Human Neocortical Neurosolver (HNN), a New Software Tool for Interpreting the Cellular and Network Origin of Human MEG/EEG Data.” eLife 9 (January). https://doi.org/10.7554/eLife.51214.

Neymotin, Samuel A., Benjamin A. Suter, Salvador Dura-Bernal, G. M. G. Shepherd, M. Migliore, and William W. Lytton. 2017. “Optimizing Computer Models of Corticospinal Neurons to Replicate in Vitro Dynamics.” Journal of Neurophysiology 117 (1): 148–62.

Nicola, Wilten, and Claudia Clopath. 2017. “Supervised Learning in Spiking Neural Networks with FORCE Training.” Nature Communications 8 (1): 2208.

O’Connell, Monica Noelle, Annamaria Barczak, Deborah Ross, Tammy McGinnis, Charles E. Schroeder, and Peter Lakatos. 2015. “Multi-Scale Entrainment of Coupled Neuronal Oscillations in Primary Auditory Cortex.” Frontiers in Human Neuroscience 9. https://doi.org/10.3389/fnhum.2015.00655.

Oliver, Douglas L., Nell B. Cant, Richard R. Fay, and Arthur N. Popper. 2018. The Mammalian Auditory Pathways: Synaptic Organization and Microcircuits. Springer.

Oswald, Manfred J., Malinda L. S. Tantirigama, Ivo Sonntag, Stephanie M. Hughes, and Ruth M. Empson. 2013. “Diversity of Layer 5 Projection Neurons in the Mouse Motor Cortex.” Frontiers in Cellular Neuroscience 7 (October): 174.

Paciello, Fabiola, Marco Rinaudo, Valentina Longo, Sara Cocco, Giulia Conforto, Anna Pisani, Maria Vittoria Podda, Anna Rita Fetoni, Gaetano Paludetti, and Claudio Grassi. 2021. “Auditory Sensory Deprivation Induced by Noise Exposure Exacerbates Cognitive Decline in a Mouse Model of Alzheimer’s Disease.” eLife 10 (October). https://doi.org/10.7554/eLife.70908.

Parasuram, Harilal, Bipin Nair, Egidio D’Angelo, Michael L. Hines, Giovanni Naldi, and Shyam Diwakar. 2016. “Computational Modeling of Single Neuron Extracellular Electric Potentials and Network Local Field Potentials Using LFPsim.” Frontiers in Computational Neuroscience 10: 65.

Park, Youngmin, and Maria N. Geffen. 2020. “A Circuit Model of Auditory Cortex.” PLoS Computational Biology 16 (7): e1008016.

Passingham, R. E. 1973. “Anatomical Differences between the Neocortex of Man and Other Primates.” Brain, Behavior and Evolution 7 (5): 337–59.

Pi, Hyun-Jae, Balázs Hangya, Duda Kvitsiani, Joshua I. Sanders, Z. Josh Huang, and Adam Kepecs. 2013. “Cortical Interneurons That Specialize in Disinhibitory Control.” Nature 503 (7477): 521–24.

Poirazi, Panayiota, and Athanasia Papoutsi. 2020. “Illuminating Dendritic Function with Computational Models.” Nature Reviews Neuroscience. https://doi.org/10.1038/s41583-020-0301-7.

Povysheva, N. V., A. V. Zaitsev, S. Kröner, O. A. Krimer, D. C. Rotaru, G. Gonzalez-Burgos, D. A. Lewis, and L. S. Krimer. 2007. “Electrophysiological Differences between Neurogliaform Cells from Monkey and Rat Prefrontal Cortex.” Journal of Neurophysiology 97 (2): 1030–39.

Prinz, Astrid A., Dirk Bucher, and Eve Marder. 2004. “Similar Network Activity from Disparate Circuit Parameters.” Nature Neuroscience 7 (12): 1345–52.

Purves, Dale, George Augustine, David Fitzpatrick, William Hall, Anthony LaMantia, Leonard White, Richard Mooney, and Michael Platt. 2018. Neuroscience.

Raveh, Eyal, Weili Hu, Blake Croll Papsin, and Vito Forte. 2002. “Congenital Conductive Hearing Loss.” The Journal of Laryngology and Otology 116 (2): 92–96.

Sakata, Shuzo, and Kenneth D. Harris. 2009. “Laminar Structure of Spontaneous and Sensory-Evoked Population Activity in Auditory Cortex.” Neuron 64 (3): 404–18.

Schroeder, Charles E. 1998. “A Spatiotemporal Profile of Visual System Activation Revealed by Current Source Density Analysis in the Awake Macaque.” Cerebral Cortex. https://doi.org/10.1093/cercor/8.7.575.

Schroeder, Charles E., Peter Lakatos, Yoshinao Kajikawa, Sarah Partan, and Aina Puce. 2008. “Neuronal Oscillations and Visual Amplification of Speech.” Trends in Cognitive Sciences 12 (3): 106–13.

Schuman, Benjamin, Robert P. Machold, Yoshiko Hashikawa, János Fuzik, Gord J. Fishell, and Bernardo Rudy. 2019. “Four Unique Interneuron Populations Reside in Neocortical Layer 1.” The Journal of Neuroscience: The Official Journal of the Society for Neuroscience 39 (1): 125–39.

Serkov, F. N., and Yu A. Gonchar. 1996. “Morphometric Characteristics of Synaptic Apparatus in the Dorsal Nucleus of the Medial Geniculate Body of the Cat.” Neurophysiology 28 (4): 155–63.

Shepherd, Gordon M. G., and Naoki Yamawaki. 2021. “Untangling the Cortico-Thalamo-Cortical Loop: Cellular Pieces of a Knotty Circuit Puzzle.” Nature Reviews. Neuroscience 22 (7): 389–406.

Sherman, Maxwell A., Shane Lee, Robert Law, Saskia Haegens, Catherine A. Thorn, Matti S. Hämäläinen, Christopher I. Moore, and Stephanie R. Jones. 2016. “Neural Mechanisms of Transient Neocortical Beta Rhythms: Converging Evidence from Humans, Computational Modeling, Monkeys, and Mice.” Proceedings of the National Academy of Sciences of the United States of America 113 (33): E4885–94.

Sivagnanam, Subhashini, Wyatt Gorman, Donald Doherty, Samuel A. Neymotin, Stephen Fang, Hermine Hovhannisyan, William W. Lytton, and Salvador Dura-Bernal. 2020. “Simulating Large-Scale Models of Brain Neuronal Circuits Using Google Cloud Platform.” In Practice and Experience in Advanced Research Computing, 505–9. PEARC ’20. New York, NY, USA: Association for Computing Machinery.

Sivagnanam, Subhashini, A. Majumdar, K. Yoshimoto, V. Astakhov, A. Bandrowski, M. E. Martone, and Nicholas T. Carnevale. 2013. “Introducing the Neuroscience Gateway.” In IWSG. Vol. 993. Citeseer.

Spencer, Kevin M., Margaret A. Niznikiewicz, Paul G. Nestor, Martha E. Shenton, and Robert W. McCarley. 2009. “Left Auditory Cortex Gamma Synchronization and Auditory Hallucination Symptoms in Schizophrenia.” BMC Neuroscience 10 (July): 85.

Spruston, Nelson. 2008. “Pyramidal Neurons: Dendritic Structure and Synaptic Integration.” Nature Reviews. Neuroscience 9 (3): 206–21.

Stanley, David A., Arnaud Y. Falchier, Benjamin R. Pittman-Polletta, Peter Lakatos, Miles A. Whittington, Charles E. Schroeder, and Nancy J. Kopell. 2019. “Flexible Reset and Entrainment of Delta Oscillations in Primate Primary Auditory Cortex: Modeling and Experiment.” BioRxiv. https://doi.org/10.1101/812024.

Steinschneider, M., D. H. Reser, Y. I. Fishman, C. E. Schroeder, and J. C. Arezzo. 1998. “Click Train Encoding in Primary Auditory Cortex of the Awake Monkey: Evidence for Two Mechanisms Subserving Pitch Perception.” The Journal of the Acoustical Society of America 104 (5): 2935–55.

Steinschneider, M., C. E. Tenke, C. E. Schroeder, D. C. Javitt, G. V. Simpson, J. C. Arezzo, and H. G. Vaughan Jr. 1992. “Cellular Generators of the Cortical Auditory Evoked Potential Initial Component.” Electroencephalography and Clinical Neurophysiology 84 (2): 196–200.

Sussillo, David, and L. F. Abbott. 2009. “Generating Coherent Patterns of Activity from Chaotic Neural Networks.” Neuron 63 (4): 544–57.

Suter, Benjamin A., Michele Migliore, and Gordon M. G. Shepherd. 2013. “Intrinsic Electrophysiology of Mouse Corticospinal Neurons: A Class-Specific Triad of Spike-Related Properties.” Cerebral Cortex 23 (8): 1965–77.

Taniwaki, T., K. Tagawa, F. Sato, and K. Iino. 2000. “Auditory Agnosia Restricted to Environmental Sounds Following Cortical Deafness and Generalized Auditory Agnosia.” Clinical Neurology and Neurosurgery 102 (3): 156–62.

Tramo, Mark Jude, Peter A. Cariani, Christine K. Koh, Nikos Makris, and Louis D. Braida. 2005. “Neurophysiology and Neuroanatomy of Pitch Perception: Auditory Cortex.” Annals of the New York Academy of Sciences 1060 (December): 148–74.

Tramo, Mark Jude, Gaurav D. Shah, and Louis D. Braida. 2002. “Functional Role of Auditory Cortex in Frequency Processing and Pitch Perception.” Journal of Neurophysiology 87 (1): 122–39.

Tremblay, Robin, Soohyun Lee, and Bernardo Rudy. 2016. “GABAergic Interneurons in the Neocortex: From Cellular Properties to Circuits.” Neuron 91 (2): 260–92.

Tripathy, Shreejoy J., Shawn D. Burton, Matthew Geramita, Richard C. Gerkin, and Nathaniel N. Urban. 2015. “Brain-Wide Analysis of Electrophysiological Diversity Yields Novel Categorization of Mammalian Neuron Types.” Journal of Neurophysiology 113 (10): 3474–89.

Turi, Gergely Farkas, Wen-Ke Li, Spyridon Chavlis, Ioanna Pandi, Justin O’Hare, James Benjamin Priestley, Andres Daniel Grosmark, et al. 2019. “Vasoactive Intestinal Polypeptide-Expressing Interneurons in the Hippocampus Support Goal-Oriented Spatial Learning.” Neuron 101 (6): 1150–65.e8.

Vijayan, Sujith, and Nancy J. Kopell. 2012. “Thalamic Model of Awake Alpha Oscillations and Implications for Stimulus Processing.” Proceedings of the National Academy of Sciences of the United States of America 109 (45): 18553–58.

Wang, Xiaoqin, Thomas Lu, Ross K. Snider, and Li Liang. 2005. “Sustained Firing in Auditory Cortex Evoked by Preferred Stimuli.” Nature 435 (7040): 341–46.

Wang, Yuan, Agnieszka Brzozowska-Prechtl, and Harvey J. Karten. 2010. “Laminar and Columnar Auditory Cortex in Avian Brain.” Proceedings of the National Academy of Sciences of the United States of America 107 (28): 12676–81.

Winer, J. A., and D. T. Larue. 1996. “Evolution of GABAergic Circuitry in the Mammalian Medial Geniculate Body.” Proceedings of the National Academy of Sciences of the United States of America 93 (7): 3083–87.

Winer, Jeffery A., Michelle L. Chernock, David T. Larue, and Steven W. Cheung. 2002. “Descending Projections to the Inferior Colliculus from the Posterior Thalamus and the Auditory Cortex in Rat, Cat, and Monkey.” Hearing Research 168 (1-2): 181–95.

Winer, Jeffery A., and Christoph E. Schreiner. 2010. The Auditory Cortex. Springer Science & Business Media.

Yamawaki, Naoki, Katharine Borges, Benjamin A. Suter, Kenneth D. Harris, and Gordon M. G. Shepherd. 2014. “A Genuine Layer 4 in Motor Cortex with Prototypical Synaptic Circuit Connectivity.” eLife 3 (December): e05422.

Yamawaki, Naoki, and Gordon M. G. Shepherd. 2015. “Synaptic Circuit Organization of Motor Corticothalamic Neurons.” The Journal of Neuroscience: The Official Journal of the Society for Neuroscience 35 (5): 2293–2307.

Zendel, Benjamin Rich, Marie-Élaine Lagrois, Nicolas Robitaille, and Isabelle Peretz. 2015. “Attending to Pitch Information Inhibits Processing of Pitch Information: The Curious Case of Amusia.” The Journal of Neuroscience: The Official Journal of the Society for Neuroscience 35 (9): 3815–24.

Zhu, J. J., W. W. Lytton, and J. T. Xue. 1999. “An Intrinsic Oscillation in Interneurons of the Rat Lateral Geniculate Nucleus.” Journal of. https://journals.physiology.org/doi/abs/10.1152/jn.1999.81.2.702.

Zhu, J. J., D. J. Uhlrich, and W. W. Lytton. 1999a. “Burst Firing in Identified Rat Geniculate Interneurons.” Neuroscience 91 (4): 1445–60.

Zhu, J. J., D. J. Uhlrich, and W. W. Lytton. 1999b. “Properties of a Hyperpolarization-Activated Cation Current in Interneurons in the Rat Lateral Geniculate Nucleus.” Neuroscience 92 (2): 445–57.

Zulfiqar, Isma, Michelle Moerel, and Elia Formisano. 2019. “Spectro-Temporal Processing in a Two-Stream Computational Model of Auditory Cortex.” Frontiers in Computational Neuroscience 13: 95.

